# Distinct ancient structural polymorphisms control heterodichogamy in walnuts and hickories

**DOI:** 10.1101/2023.12.23.573205

**Authors:** Jeffrey S. Groh, Diane C. Vik, Kristian A. Stevens, Patrick J. Brown, Charles H. Langley, Graham Coop

## Abstract

The maintenance of stable mating type polymorphisms is a classic example of balancing selection, underlying the nearly ubiquitous 50/50 sex ratio in species with separate sexes. One lesser known but intriguing example of a balanced mating polymorphism in angiosperms is heterodichogamy – polymorphism for opposing directions of dichogamy (temporal separation of male and female function in hermaphrodites) within a flowering season. This mating system is common throughout Juglandaceae, the family that includes globally important and iconic nut and timber crops – walnuts (*Juglans*), as well as pecan and other hickories (*Carya*). In both genera, heterodichogamy is controlled by a single dominant allele. We fine-map the locus in each genus, and find two ancient (*>*50 Mya) structural variants involving different genes that both segregate as genus-wide trans-species polymorphisms. The *Juglans* locus maps to a ca. 20 kb structural variant adjacent to a probable trehalose phosphate phosphatase (*TPPD-1*), homologs of which regulate floral development in model systems. *TPPD-1* is differentially expressed between morphs in developing male flowers, with increased allele-specific expression of the dominant haplotype copy. Across species, the dominant haplotype contains a tandem array of duplicated sequence motifs, part of which is an inverted copy of the *TPPD-1* 3’ UTR. These repeats generate various distinct small RNAs matching sequences within the 3’ UTR and further downstream. In contrast to the single-gene *Juglans* locus, the *Carya* heterodichogamy locus maps to a ca. 200-450 kb cluster of tightly linked polymorphisms across 20 genes, some of which have known roles in flowering and are differentially expressed between morphs in developing flowers. The dominant haplotype in pecan, which is nearly always heterozygous and appears to rarely recombine, shows markedly reduced genetic diversity and is over twice as long as its recessive counterpart due to accumulation of various types of transposable elements. We did not detect either genetic system in other heterodichogamous genera within Juglandaceae, suggesting that additional genetic systems for heterodichogamy may yet remain undiscovered.

## Introduction

Flowering plants have evolved a remarkable diversity of sexual heteromorphisms (e.g. dioecy, heterostyly, gynodioecy, and mirror-image flowers) that have long fascinated evolutionary biologists (Darwin 1877; Charlesworth and Charlesworth 1979; Barrett 2010). Such systems present key opportunities to test evolutionary theories of sexual reproduction and to understand its ecological and genetic consequences. Heterodichogamy presents an intriguing and relatively under-explored example of a balanced polymorphism in the sexual organization of some angiosperms, in this case not in space but in time. In Darwin’s words, heterodichogamous populations *“consist of two bodies of individuals, with their flowers differing in function, though not in structure; for certain individuals mature their pollen before the female flowers on the same plant are ready for fertilisation and are called proterandrous* [protandrous]*; whilst conversely other individuals, called proterogynous* [protogynous]*, have their stigmas mature before their pollen is ready”*(1877). This system generates strong disassortative mating between morphs (Bai *et al*. 2007), thus classical sex ratio theory predicts a 50/50 ratio of protandrous and protogynous individuals at equilibrium (Gleeson 1982). Although heterodichogamy has likely evolved at least a dozen times in angiosperms (Renner 2001; Endress 2020), to our knowledge the genetic loci controlling the inherited basis of this mating system have not been described at the molecular level in any species. Thus, it is not known whether similar genetic mechanisms control heterodichogamy in different taxa, nor whether the loci involved experience similar evolutionary dynamics as sex chromosomes and other supergenes that underpin other complex mating polymorphisms (Charlesworth 2016; Gutíerrez-Valencia *et al*. 2021).

Heterodichogamy is well-known within Juglandaceae (Delpino 1874; Darwin 1876; Pringle 1879; Stuckey 1915, Fig. 1A,B), the family that includes two major globally important nut and timber-producing groups of trees – the walnuts (*Juglans*) and hickories (*Carya*, includes cultivated pecans). The ancestor of Juglandaceae evolved unisexual, wind-pollinated flowers from an ancestor with bisexual flowers near the Cretaceous–Paleogene boundary (Friis 1983; Sims *et al*. 1999), and a rich fossil history shows a radiation of Juglandaceae during the Paleocene (Manchester 1987, 1989). In both walnuts and hickories, protogyny is inherited via a dominant Mendelian allele (dominant allele – *G*, recessive allele – *g*, Gleeson 1982; Thompson and Romberg 1985). Other genera phylogenetically closer to *Juglans* (*Cyclocarya, Platycarya*) are also known to be heterodichogamous (Fukuhara and Tokumaru 2014; Mao *et al*. 2019). These observations have suggested a single origin of the alleles controlling heterodichogamy in the common ancestor of these taxa. Here, we identify and characterize the evolutionary history of the genetic loci controlling heterodichogamy in both *Juglans* and *Carya*. We find two distinct and substantively different genetic underpinnings for heterodichogamy in these genera, and show that each genetic system is an ancient trans-species balanced polymorphism for a structural variant.

**Figure 1:**
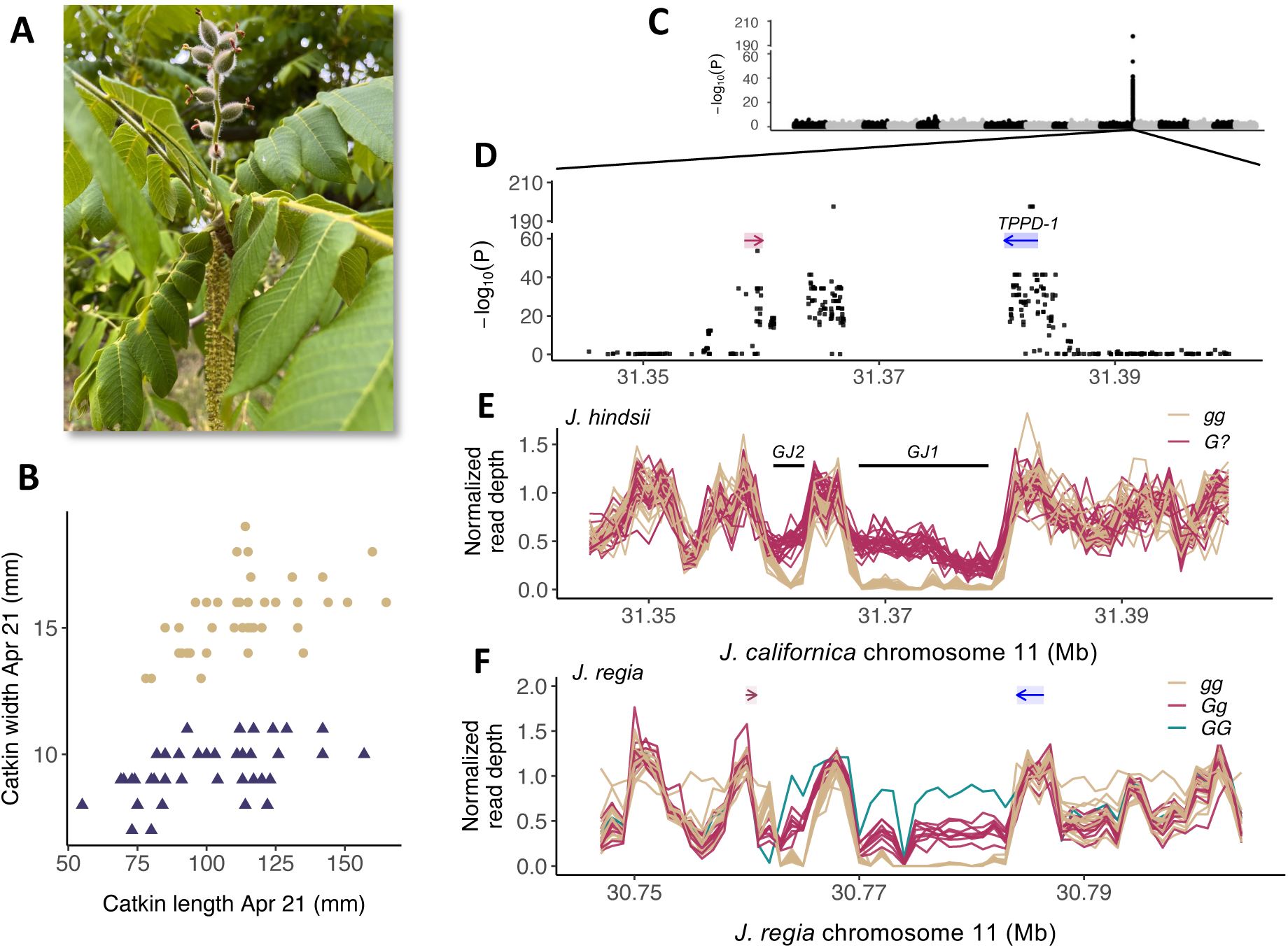
**A**) A protogynous individual of *J. ailantifolia* with developing fruits and catkins that are shedding pollen. Photo by JSG. **B)** Protandrous (tan) and protogynous (dark blue) morphs of *J. hindsii* are readily distinguished by catkin size in the first half of the flowering season. **C)** GWAS of flowering type in 44 individuals of *J. hindsii* from a natural population, against a long-read assembly of the sister species, *J. californica*. **D)** The GWAS peak occurs across and in the 3’ region of *TPPD-1*, a probable trehalose-6-phosphate phosphatase gene (blue rectangle, arrow indicates direction of transcription). Red bar shows the position of a *NDR1/HIN1* –like gene. **E)** Normalized average read depth in 1 kb windows for 44 *J. hindsii* from a natural population against a genome assembly for *J. californica*. Black bars show positions of identified indels. **F)**Normalized average read depth in 1 kb windows for 26 *J. regia* individuals in the region syntenic with the *J. hindsii* G-locus. Colored bars and arrows indicate positions of orthologous genes bordering the G-locus.

### Ancient regulatory divergence controls heterodichogamy across *Juglans*

In a natural population of Northern California black walnut (*J. hindsii*), protandrous and protogynous morphs were found to occur in roughly equal proportions (43 vs. 38, P=0.47 under null hypothesis of equal proportions). A genome-wide association study (GWAS) for dichogamy type in this population recovered a single, strong association peak, consistent with the known Mendelian inheritance of heterodichogamy type in *Juglans* (Fig. 1C, Gleeson 1982). This region is syntenic with the top GWAS hit location for dichogamy type in *J. regia* (Fig. S1, Bernard *et al*. 2020) as well as a broad region associated with dichogamy type in *J. nigra* (Chatwin *et al*. 2023). The association peak extends roughly 20 kb (hereafter, the G-locus) across a probable trehalose-6-phosphate phosphatase gene (*TPPD-1*, Fig. 1D, corresponding to LOC108984907 in the Walnut 2.0 assembly). TPPs catalyze the dephosphorylation of trehalose-6-phosphate to trehalose as part of a signaling pathway essential for establishing regular flowering time in Arabidopsis (Wahl *et al*. 2013) and inflorescence architecture in maize (Satoh-Nagasawa *et al*. 2006). *TPP* genes were recently identified as differentially expressed across the transition to flowering in both *J. sigillata* and *J. mandshurica* (Lu *et al*. 2020; Li *et al*. 2022b). In *J. regia*, *TPPD-1* is expressed in a broad range of vegetative and floral tissues (Fig. S2), and we detected full-length transcripts from multiple tissues related to flowering in *J. regia* homozygotes for both G-locus haplotypes (Table S2). Consistent with the width of the GWAS hit, we observed strong linkage disequilibrium (LD) extending across this region, suggesting locally reduced recombination in the genealogical history of the sample for both *J. hindsii* and *J. regia* (Fig. S3).

We identified copy number variation consistent with three *>*1 kb indels distinguishing G-locus haplotypes (*GJ1* and *GJ2* present in the *G* haplotype; *gJ3* in the *g* haplotype, Figs. 1E, S1). Consistent with the known dominance, all *J. hindsii* protogynous individuals (*G* ?) appear to be hemizygous for these indels, while protandrous individuals (*gg*) are homozygous for *gJ3*. The apparent absence of any homozygotes for the *G* haplotype in our *J. hindsii* sample is consistent with disassortative mating at the G-locus in a natural population (P=0.059). Strikingly similar copy number patterns were seen in *J. regia* (Fig. 1F, S1), with a known homozygote for the protogynous allele (‘Sharkey’, *GG*, Gleeson 1982) lending further support for these indels. To look for the presence of this structural variant across *Juglans* species, we generated whole-genome resequencing data from both morphs from 5 additional *Juglans* species (Table S1) combined with existing data from known morphs in two additional species (Stevens *et al*. 2018). Parallel patterns of read depth at the G-locus indicate that the same structural variant segregates in perfect association with dichogamy type across all nine species examined (Fig. 1F, S4). As our species sampling spans the deepest divergence event in the genus (Manchester 1987; Mu *et al*. 2020), this suggested that the G-locus haplotypes may have diverged in the common ancestor of *Juglans*.

Consistent with the *Juglans* G-locus being an ancient balanced polymorphism, nucleotide divergence between G-locus haplotypes is exceptionally high compared to the genome-wide background (falling in the top 1% of values across the entire aligned chromosome in several species, Fig. 2D), and is comparable to the divergence observed between *Juglans* and *Carya* in the same region (Fig. S6). Applying a molecular clock, we estimated the age of the G-locus haplotypes to be 41-69 Mya (methods). As both fossil and molecular evidence place the most recent common ancestor of *Juglans* in the Eocene (Manchester 1987; Mu *et al*. 2020; Zhou *et al*. 2021), this estimate is consistent with G-locus haplotypes originating in the common ancestor of extant *Juglans*.

**Figure 2:**
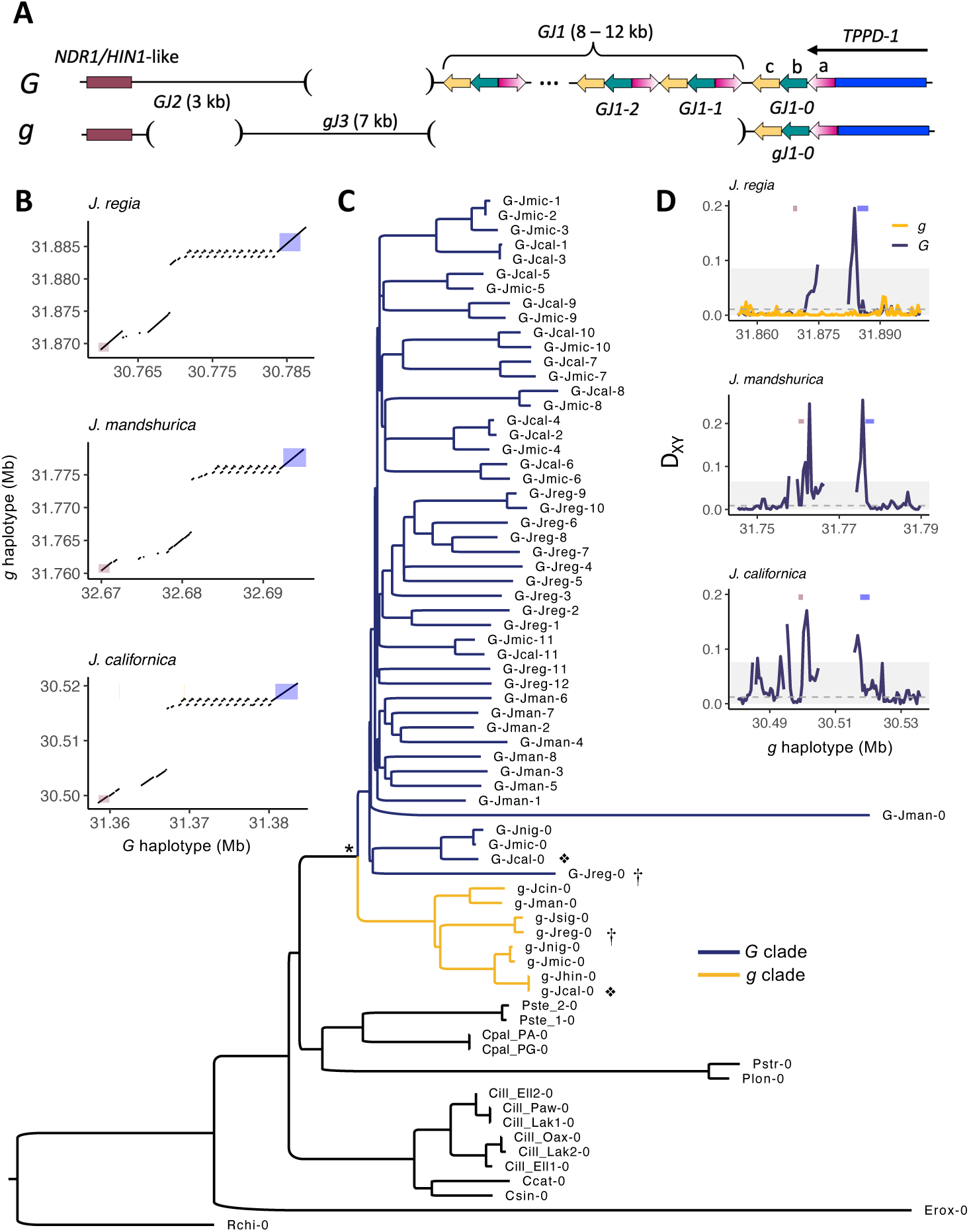
**A**) Schematic of the G-locus structural variant in *Juglans*. Presence/absence of indels are indicated by continuous lines vs. open parentheses. Maroon and blue bars show bordering genes. Black arrow across *TPPD-1* indicates direction of transcription. Colored arrows represent subunits of the repeat motif within the *GJ1* indel and paralogous sequence outside of the indel. Each repeat motif (numbered 1,2,…) is comprised of subunits *a*, *b*, and *c*. Subunit *a*, which is homologous with the 3’ UTR of *TPPD-1*, is inverted within the indel. **B)** Dotplots showing pairwise alignment of alternate haplotypes for three species representing the major clades within the genus. Maroon and blue rectangles indicate the locations of the genes bordering the G-locus as in (A). **C)** Phylogeny of GJ1 repeats. Sample codes from L-R indicate: haplotype (for *Juglans*only), abbreviated binomial, optional additional identifier, and repeat number (as in A) (see Table S3 for full taxon list and data sources). Sequences from *g* haplotypes (gold) and *G* haplotypes (blue) form two sister clades (**\***, 99% bootstrap support) whose most recent common ancestor predates the radiation of *Juglans*. Daggers and diamonds highlight representative sequences from *J. californica* and *J. regia* that show the trans-species polymorphism. **D)** Average nucleotide divergence in 500 bp windows between alternate G-locus haplotypes for three within-species comparisons. Dotted gray lines indicate the chromosome-wide average, and shaded gray regions indicate 95% quantiles. Maroon and blue bars indicate positions of genes as in (A).

We next examined variation within the *TPPD-1* gene sequence. In a comparison of both haplotypes from three species spanning the earliest divergence events within *Juglans*, we find several genus-wide transspecific polymorphic SNPs within the 3’ UTR of *TPPD-1*, but none within coding sequence, indicating recombination over deep timescales and a lack of shared functional coding divergence (Fig. S5B). However, we see numerous coding polymorphisms that are fixed differences between haplotypes within species and that are shared across more closely related species (Fig. S5C), suggesting the intriguing possibility of recruitment and turnover of functional divergence within the *TPPD-1* gene sequence.

We next investigated the three indels between the *G* and *g* haplotypes (Fig. 2A). The indels *GJ2* and *gJ3* are not obvious functional candidates. For example, *gJ3* is derived from an insertion of a CACTA-like DNA transposon in *g* haplotypes of *Juglans*, which lacks evidence of gene expression or strong sequence conservation across species (Fig. S7). Nonetheless, given their conserved presence it seems plausible that these indels have some role in the establishment of dichogamy types.

Turning to the *GJ1* indel closer to *TPPD-1*, using pairwise alignments we found that *GJ1* contains a series of *∼*1kb tandem repeats that are paralogous with sequence in the 3’ region of *TPPD-1* (Fig. 2A,B). We find no evidence for the inclusion of the repeat in full length *TPPD-1* transcripts from a *GG J. regia* individual. Each repeat, ordered *GJ1-1* up to *GJ1-12* moving downstream of *TPPD-1*, consists of three subunits. One subunit (*GJ1a*) is an inverted *∼*250-300bp motif homologous with the 3’ UTR of *TPPD-1*. The other two subunits (*GJ1b* and *GJ1c*) are non-inverted *∼*300bp motifs homologous with regions within 1kb downstream of the 3’ UTR. These tandem duplicates are present in *G* haplotypes across multiple species with long read assemblies, varying in number from 8-12 copies (Fig. 2B). Consistent with G-locus haplotypes having arisen in the ancestor of *Juglans*, a maximum-likelihood phylogeny of concatenated aligned sequence from *GJ1* repeats and their homologous sequence immediately downstream of the *TPPD-1* (Table S3) shows that all *G* haplotype sequences form a sister clade to all *g* haplotype sequences (Fig. 2C). Among more closely related species (*J. microcarpa* and *J. californica*, *∼*5-10 Mya species divergence) we found evidence of conservation of specific repeats, with some repeats from the same relative positions in different species clustering together in the phylogeny. On the other hand, for deeper divergences (*∼*40-50 Mya species divergence), evolution of *GJ1* repeats is characterized by lineage-specific turnover due to expansion and contraction of the repeat array and/or possible gene conversion. For example, the majority of repeats within *J. regia* are most closely related to repeats at other positions within the same species, and likewise for *J. mandshurica*.

The developmental basis of heterodichogamy in *J. regia* is driven by a differential between protogynous and protandrous morphs in the extent of both male and female floral primordia differentiation in the growing season prior to flowering (Luza and Polito 1988; Polito and Pinney 1997). Correspondingly, we found that protandrous and protogynous morphs of *J. regia* differ in the size of male catkin buds by mid-summer (Fig. S9). Given the conserved non-coding *G* haplotype indels, and the lack of *Juglans*-wide trans-specific coding polymorphism in *TPPD-1*, we hypothesized that the developmental differential between dichogamy types may be due to differential regulation of *TPPD-1* during early floral development. We reanalyzed RNAseq data collected from male and female flower buds of *J. mandshurica*, where a *TPP* gene was one of a number of differentially expressed genes over the course of flowering (Li *et al*. 2022b). Across two separate *J. mandshurica* datasets (Qin *et al*. 2021; Li *et al*. 2022b), the *TPPD-1* ortholog shows increased expression in male buds from samples with protogynous genotypes (Figs. S10, S11). Allelic depth at G-locus SNPs in these data indicated that this is driven by higher allele-specific expression of the *G* haplotype *TPPD-1* (Figs. S10, S11). We found a parallel bias in allele expression in transcriptomic data from multiple tissues of a single individual of *J. regia* that is heterozygous at the G-locus (Fig. S12) (Dang *et al*. 2016). To explore possible mechanisms by which variation at the G-locus might regulate *TPPD-1* expression, we screened publicly available small RNA sequence libraries from male and female floral buds of a protandrous and protogynous morph in *J. mandshurica* (Li *et al*. 2023). We found numerous 18-24 bp small RNAs in protogynous male buds (fewer in female buds and none in a protandrous sample) that map to various repeats within the *GJ1* indel and downstream of the coding region of *TPPD-1* (Figs. S13, S14). The most abundant of these small RNAs maps to the *GJ1c* subunit, at a conserved site among repeats across species, with the small RNA closely matching a *G* haplotype sequence just downstream of the 3’ UTR that is absent from *g* haplotypes (Fig. S15). A number of the other distinct small RNAs sequences show perfect sequence matching with the 3’ UTR of *TPPD-1*; some matching both G-locus alleles, and some that match only *G* alleles (Fig. S16).

Taken together, multiple independent lines of evidence support that the control of dichogamy type in *Juglans* is governed by *TPPD-1* regulatory variation that predates the radiation of the genus: (1) the genus-wide balanced polymorphism of the array of (*≥*8) 3’ UTR homologous repeats, (2) the production of small RNAs by this *G* haplotype repeat unit matching sites within and downstream of the 3’ UTR, (3) and differences in *TPPD-1* expression between the morphs due to higher allele-specific expression of the *G* haplotype *TPPD-1*. (4) Finally, the presumed substrate of *TPPD-1*, T6P, has been found in model systems to play a critical role in regulating the transition to flowering. In Arabidopsis for instance, overexpression of a heterologous *TPP* delays flowering (Schluepmann *et al*. 2003), as does knock down of a T6P synthase gene (Wahl *et al*. 2013). These facts are consistent with the idea that higher expression of *TPPD-1* in developing male flowers of protogynous walnuts could be responsible for delayed male flowering. While the mechanistic basis of the expression difference is not clear, we note that some small RNA pathways upregulate gene expression (Li *et al*. 2006; Shibuya *et al*. 2009; Fröhlich and Vogel 2009). Further investigation of the functional role of these RNAs is clearly warranted, given an emerging view of the general importance of small RNAs in plant sex-determining systems (Akagi *et al*. 2014; Müller *et al*. 2020).

Finally, we note that while the divergence of G-locus haplotypes predates the *Juglans* radiation, our phylogeny inference suggests it is more recent than the divergence of *Juglans* with its closest cousins, *Pterocarya* and *Cyclocarya*, and the relationships of non-*Juglans* sequences resemble previously estimated species trees for these taxa (Fig. 2C). Consistent with this, we found no evidence of the G-locus structural variant in resequencing data from multiple individuals from each of *Cyclocarya*, *Pterocarya*, and *Platycarya* (Fig. S8).

### A supergene controls heterodichogamy in *Carya*

While heterodichogamy has been suggested as the ancestral state of Juglandaceae, our results indicate that the *Juglans* G-locus is specific to walnuts. To identify the basis of the trait in pecan (*C. illinoinensis*), we generated whole-genome resequencing data from 18 pecan varieties of known dichogamy type and combined these with existing data for 12 additional varieties (Xiao *et al*. 2021) to perform a GWAS for dichogamy type. We identified a single peak of strong association at a locus on chromosome 4, with many alleles in strong LD (Fig. 3, S17), consistent with previous QTL mapping (Bentley *et al*. 2019). This pecan G-locus region is not homologous with the *Juglans* G-locus, implying either convergent origins of heterodichogamy within Juglandaceae, or a single origin followed by turnover in the underlying genetic mechanism.

**Figure 3:**
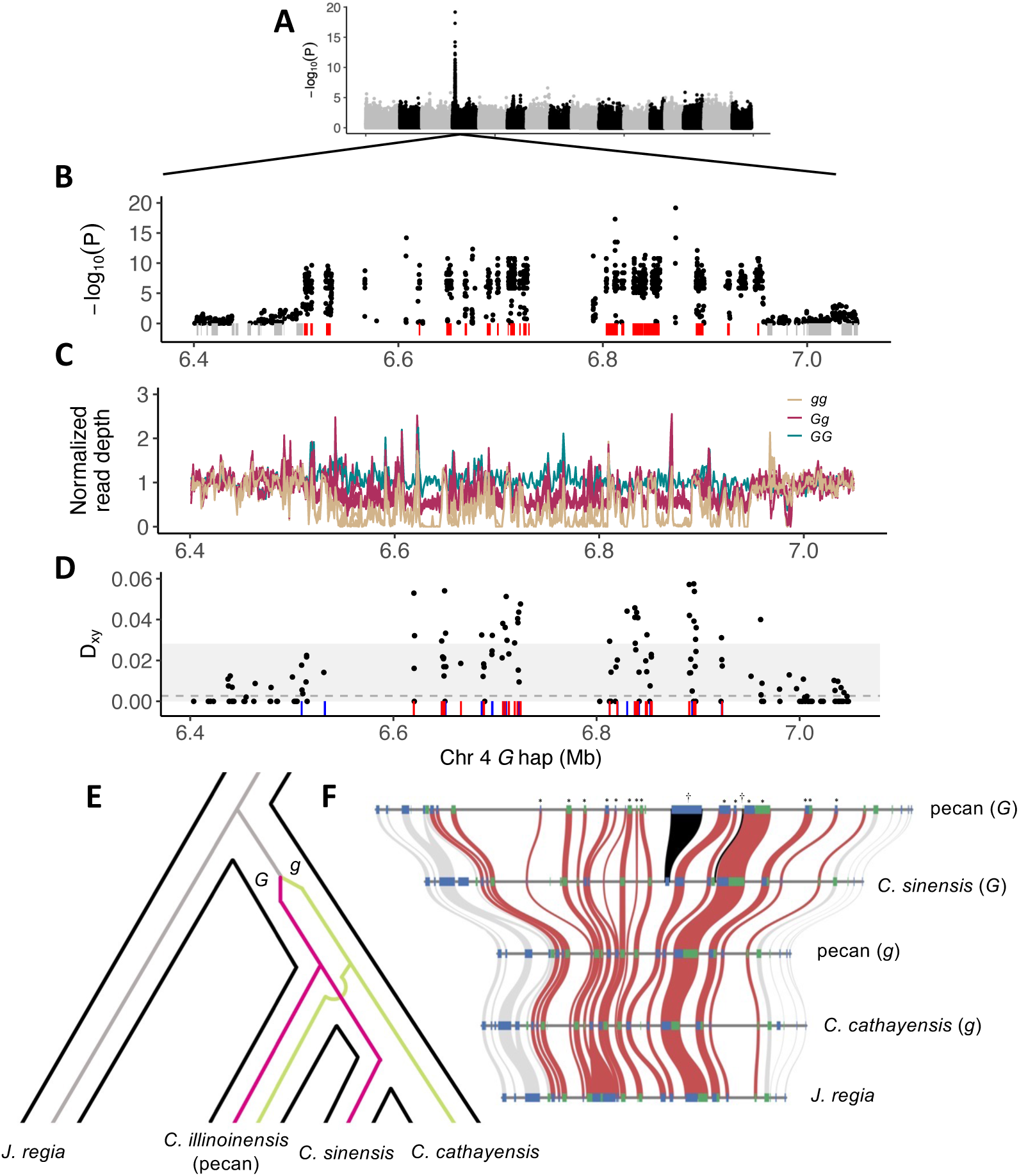
The G-locus in *Carya*. **A**) GWAS for dichogamy type in 30 pecan varieties identifies a single peak of strong association on chromosome 4. **B)** Zoomed in view of the GWAS peak. Boxes underneath plot indicate positions of predicted genes. Red boxes are those that fall within the region of strong association. **C)** Normalized average read depth in 1 kb windows across the location of the GWAS hit reveals a structural variant segregating in perfect association with dichogamy type. **D)** Nucleotide divergence between G-locus haplotypes within coding sequence is strongly elevated against the genome-wide background. Dotted line shows the genome-wide average for coding sequence, shaded interval shows 99% quantile. Colored tick marks at bottom show locations where other *Carya* species are heterozygous for SNPs that are fixed between pecan G-locus haplotypes. Blue – North American, red – East Asian. **E)** The phylogeny of the G-locus is discordant with the species tree, reflecting trans-species polymorphism. *Carya* G-locus haplotypes diverged after the split with *Juglans* and were present in the most recent common ancestor of *Carya*. **F)** Gene-level synteny for assemblies of both G-locus haplotypes in pecan and East Asian *Carya*, and *J. regia*. *G* haplotypes are longer than *g*haplotypes in both major *Carya* clades due to accumulation of transposable elements. Colored boxes show positions of genes in each assembly, with color indicating the strand (blue – plus, green – minus). Red bands connect orthologous genes that fall within the GWAS peak in pecan; gray bands connect genes outside the region. Individual gene trees that support the trans-species polymorphism are indicated with asterisks along the top. Black bands and daggers indicate two genes that appear to be uniquely shared by North American and East Asian *G* haplotypes.

We identified a large segregating structural variant in pecan (445 kb in the primary haplotype-resolved assembly of ‘Lakota’, *Gg* diploid genotype) overlapping the location of the GWAS peak that is perfectly associated with dichogamy type (Fig. 3C), and confirmed the genotype of two known protogynous allele homozygotes (*GG*), ‘Mahan’ and ‘Apache’ (Thompson and Romberg 1985; Bentley *et al*. 2019). Coverage patterns indicated the primary assembly of ‘Lakota’ is of the *G* haplotype. Consistent with this, coverage patterns against the alternate assembly of ‘Lakota’ and against the reference assembly of ‘Pawnee’ (protandrous, *gg*) identified both as *g* haplotype assemblies (Fig. S20).

In contrast to the *Juglans* G-locus, the *Carya* G-locus contains approximately 20 predicted protein-coding genes (Fig. 3B, red boxes, Table S4). Gene-level synteny is highly conserved between the *Carya* G-locus haplotypes (Fig. 3F). Segregating copy number variation is largely localized to intergenic regions, though we see several indels within coding sequence (Fig. S18). Several of these genes have well-characterized homologs involved in flowering (e.g. *FIL1, EMS1, SLK2, CEN*, see Table S4) with plausible roles in the development of alternate dichogamy types. We found 237 nonsynonymous fixed differences between the pecan haplotypes across these 20 genes, suggesting many potential functional candidates. However, applying the McDonald-Kreitman test to each gene, the ratio of nonsynonymous to synonymous substitutions vs. polymorphisms did not identify any signals of adaptive evolution. We note however that two of these genes with annotated functional roles in flowering, *EMS1* and *FIL1*, were previously identified in a small set of highly differentially expressed genes between male catkin buds of protandrous and protogynous cultivars in the season prior to bloom near when a differential in anther development is first established (Rhein *et al*. 2023). Arabidopsis mutants of *EMS1* fail to form pollen tetrads (Zhao *et al*. 2002), and protogynous pecans have a delayed progression from pollen tetrads to mature pollen as compared to protandrous pecans (Stuckey 1915). *FIL1* homologs are expressed uniquely or predominantly in stamens (Nacken *et al*.1991, Fig. S19) and were found to be a downstream target of the B class MADS-box transcription factor *DEFICIENS* in Antirrhinum (Nacken *et al*. 1991) (the ortholog of Arabidopsis *AP3*).

We found strongly elevated nucleotide divergence between *Carya* G-locus haplotypes against the genomewide background, suggesting that they could have been maintained as an ancient balanced polymorphism (Fig. 3D). To search for evidence of trans-species polymorphism of the *Carya* G-locus, we compared the North American pecan to two long-read genome assemblies from species in the East Asian *Carya* clade (*C. sinensis* and *C. cathayensis*, Zhang *et al*. 2023), spanning the deepest split within the genus (see Fig. 3E for species tree). Patterns of read depth of the pecan whole-genome resequencing reads against these two assemblies suggested that the *C. cathayensis* and *C. sinensis* assemblies represent the *g* and *G* haplotypes, respectively (Fig. S21). This was further supported by elevated nucleotide divergence between the assemblies in this region (Fig. S22) and shared nonsynonymous coding polymorphisms with pecan haplotypes (Fig. S18). We next constructed a maximum-likelihood consensus phylogeny from concatenated protein-coding sequence from the 20 G-locus genes, and found that it conflicts with the species tree, with *C. cathayensis* clustering with the pecan *g* haplotype and *C. sinensis* clustering with the pecan *G* haplotype (both 100% bootstrap support). Finally, we used whole-genome resequencing data from 16 individuals (15 from Huang *et al*. 2019, 1 from this study) representing 15 different *Carya* species to examine heterozygosity at the G-locus across the genus. We ascertained in pecan a set of 454 SNPs in coding sequence that were fixed between G-locus haplotypes. We found 8/16 individuals in this sample that are highly heterozygous at these SNPs (25-74%, the other half being 1-5%), consistent with these individuals being heterozygous for the G-locus and with dichogamy types being maintained at 50/50 proportions in species across the entire genus (Figs. S23, S24). We conclude from these multiple lines of evidence that the *Carya* G-locus haplotypes are at least as old as the divergence between Eastern North American and East Asian clades of *Carya*, a depth of *∼*25 Mya (Zhang *et al*. 2013; Mu *et al*. 2020; Zhou *et al*. 2021).

If variation at the *Carya* G-locus controlled dichogamy types in the ancestor of *Juglans* and *Carya*, we would expect that nucleotide divergence between the *Carya* G-locus haplotypes should match nucleotide divergence between pecan and the orthologous sequence in *Juglans*. Divergence between *Carya* G-locus haplotypes at four-fold degenerate sites across the core set of *Carya* G-locus genes that group by haplotype was significantly less than that between either *Carya* haplotype and *J. regia* (0.055 vs. 0.067, P *<* 0.001), implying that the *Carya* G-locus haplotypes diverged more recently than the *Juglans*-*Carya* split. Assuming a molecular clock and a divergence time of 58-72 Mya between *Juglans* and *Carya* (Zhang *et al*. 2013; Mu *et al*. 2020; Zhou *et al*. 2021), we estimate the age of the *Carya* G-locus haplotypes to be 48-59 Mya.

The *G* haplotype is rarely found in homozygotes due to strong disassortative mating under heterodichogamy, and it seems to experience little recombination (Fig. S17), so it may experience similar evolutionary forces to non-recombination portions of Y chromosomes (Bachtrog 2013). The pecan *G* haplotype (445 kb) is over twice as large as the *g* haplotype (200 kb), largely due to a difference in transposable element content within the region of reduced recombination (*g*: 40%, *G*: 65%, genome average: 53%, Fig. S25). Part of the transposable element expansion in the *G* haplotype may be ancient, given parallel coverage patterns and assembly lengths for the East Asian G-locus haplotypes; the *C. sinensis G* assembly is *∼*350 kb, while the *C. cathayensis g* assembly is *∼*230 kb. This proliferation of transposable elements within *G* haplotypes might reflect reduced efficacy of selection on the *G* haplotype due to its lower effective population size and the enhanced effects of linked selection due to its low recombination rate (Charlesworth and Charlesworth 2000). Consistent with *G* haplotypes having a reduced effective population size and/or stronger linked selection, we observe a *∼*6 fold reduction in genetic diversity in coding regions for *G* haplotypes compared to *g* haplotypes (*π* = 0.0111 vs. 0.0018, P *<* 0.013).

Theory and data suggests that supergenes (e.g. sex chromosomes) may progressively assemble over time, perhaps through the accumulation of morph-antagonistic variation. While broad synteny of 20 predicted protein-coding genes across *Carya* G-locus assemblies is conserved as far back as the divergence with oaks (*∼*90 Mya, Larson-Johnson 2016) (Fig. S26), we do note several predicted protein-coding genes that differ among *Carya* G-locus assemblies (Fig. 3F, Table S5). Notably, two of these appear to be derived and shared between North American and East Asian *G* haplotypes and so warrant further attention (Table S5, methods). While the majority of gene trees across the G-locus support the trans-species polymorphism for the four *Carya* assemblies, we also note that the three genes at the left border (including *EMS1*) and the gene at the right border instead support the species phylogeny for these four taxa (Fig. 3F). Furthermore, we do not see clear evidence for trans-species polymorphism within these genes in the East Asian clade (Figs. 3D, S24). These observations, and the lower divergence between pecan G-locus halotypes in the three genes at the left border, suggest the boundary of recombination suppression at the G-locus may have changed since the divergence of the North American and East Asian *Carya*, perhaps after the appearance of genetic variation in neighboring genes with antagonistic fitness effects between the two dichogamy morphs.

### Genetic systems for heterodichogamy in other genera

Our estimate of the age of the *Carya* G-locus haplotypes suggests that we do not expect to find these haplotypes segregating in other known heterodichogamous genera within Juglandaceae – *Cyclocarya*, *Platycarya*, and likely *Pterocarya* – which are all more closely related to *Juglans* than to *Carya*. Nonetheless, we considered the possibility that a subset of the pecan G-locus variation could be ancestral across heterodichogamous genera, or that there has been convergent use of the same region. Therefore, we examined heterozygosity at the region syntenic with the *Carya* G-locus in a sample of 13 *Pterocarya*, 12 diploid *Cyclocarya* (including protandrous and protogynous individuals, Qu *et al*. 2023), and 3 *Platycarya strobilaceae*, but we found no evidence of increased polymorphism in this region comparable to the patterns seen in *Carya* (Fig. S27). As our analyses also indicate that heterodichogamy in these genera is not controlled by the *Juglans* G-locus (Fig. 2C, S8, S28), this suggests the intriguing possibility that additional genetic systems controlling heterodichogamy in other Juglandaceae genera remain undiscovered.

## Discussion

The genetic mechanisms underlying heterodichogamy appear to be more diverse than previously appreciated. Within Juglandaceae, two distinct ancient structural variants underlie this mating polymorphism in different genera. The identification of these loci opens further opportunities to study the ecology and the molecular and cellular mechanisms of this mating system with potential agricultural benefits in these important crop species. Our data cannot rule out the possibility that these genetic systems originated independently through the convergent evolution of heterodichogamy. However, given the clade-wide presence of heterodichogamy across Juglandaceae, it seems plausible that a heterodichogamous mating system evolved once in the ancestor and that genetic control of dichogamy type has been subject to turnover. Similar dynamics have been discovered in sex determination systems as new data and experiments improve both the phylogenetic scale and resolution of sex-chromosome turnover (Myosho *et al*. 2012; Jeffries *et al*. 2018; Hu *et al*. 2023). As heterodichogamy unites concepts of inbreeding avoidance, sexual interference, and sex-ratio selection, these systems may offer an important complement for testing theories on the evolution of the control and turnover of sexual systems (Sargent *et al*. 2006; Van Doorn and Kirkpatrick 2007; Kozielska *et al*. 2010; Blaser *et al*. 2013; Saunders *et al*. 2018). In summary, the evolution of heterodichogamy within Juglandaceae showcases both dynamic evolution and remarkable stability.

## Acknowledgements

We are grateful to UC Davis Putah Creek Riparian Reserve, Gene Cripe of Linwood Nursery, CA, USDA Wolfskill Experimental Orchard, Sonoma Botanical Garden, and UC Botanical Garden at Berkeley for access to leaf tissue from natural and cultivated collections of Juglandaceae. We thank Chuck Leslie for providing phenotype records for trees from the USDA germplasm database. We thank Jeff Ross-Ibarra, Quentin Cronk, the Dandekar lab at UC Davis, and the Coop lab for helpful discussions. Funding was provided by the United States Department of Agriculture (NIFA SCRI-Award no. 2012-51181-20027 awarded to PJB, CHL), the National Institutes of Health (NIH R35 GM136290 awarded to GC), and the National Science Foundation (NSF DISES 2307175 to GC, and NSF 1650042 awarded to JSG).

## Materials and Methods

### Phenotyping

We phenotyped 81 individuals of a naturally-occuring population of *J. hindsii* from the UC Davis Putah Creek Riparian Reserve, along a *∼*2 mile creek-side path, in spring of 2022 and 2023. Dichogamy type was ascertained visually based on the relative developmental stages of male and female flowers. To validate our assessment of flowering phenotype we measured the length and width of 1-3 catkins of each tree on the same day and plotted the size distribution of catkins in the sample (Fig. 1A). We similarly obtained dichogamy phenotypes for individuals of *J. ailantifolia*, *J. californica*, *J. cathayensis*, *J. cinerea*, and *J. major* from trees at USDA Wolfskill Experimental Orchard in spring 2023. Phenotypes of newly sequenced *C. illinoinensis* from UC Davis orchards were also scored. Dichogamy phenotypes were available for 26 previously sequenced individuals of *J. regia* from UC Davis walnut breeding program records (Stevens *et al*. 2018). Flowering phenotypes for *J. microcarpa* and *J. hindsii* trees that were previously sequenced in Stevens *et al*. (2018) were obtained from the USDA Wolfskill Experimental Orchard database (Chuck Leslie, personal communication). Phenotypes of previously sequenced *J. nigra* individuals (Stevens *et al*. 2018) were obtained from Chatwin *et al*. (2023). Phenotypes of previously sequenced varieties of pecan (Xiao *et al*. 2021) were obtained from Bentley *et al*. (2019). Phenotypes of newly sequenced pecan varieties were scored visually in spring of 2023 or otherwise obtained from Bentley *et al*. (2019) and the USDA database (https://cgru.usda.gov/carya/pecans/).

We measured the length of dormant catkin buds in 30 *J. regia* individuals on July 20, 2023. Measurements were made blind with respect to phenotype with the exception of 3 varieties which were known a priori. We took an average measurement of 6-12 of the largest easily accessible catkins using hand calipers. We tested for a difference in length using a linear model with leafing date as a covariate (Fig. S9).

### Genomic sequencing and data curation

We generated whole-genome resequencing (mean coverage *∼*35x) data for 46 *J. hindsii*; 2 individuals each from *J. ailantifolia*, *J. californica*, *J. cathayensis*, *J. cinerea*, *J. major*; 19 *C. illinoinensis*; 1 *C. ovata*; 13 *Pterocarya stenoptera*; 2 *Pterocarya rhoifolia*; 1 *Pterocarya macroptera*; and 2 *Platycarya strobilaceae* (Table S1). Samples of *J. hindsii* were a subset of trees phenotyped from the UC Davis Putah Creek Riparian Reserve, CA, plus two additional trees without dichogamy phenotypes from the same population. Other sequenced *Juglans* trees were from USDA Wolfskill Experimental Orchard. Samples of *Pterocarya stenoptera* were obtained from Wolfskill and UC Davis campus. Samples of *Carya illinoinensis* were obtained from UC Davis orchards and the original orchard of Linwood Nursery in Turlock, California. Leaf tissue was flash frozen in liquid nitrogen and preserved at –80 degrees Celsius. DNA was extracted using the Qiagen DNeasy Plant Pro Kit.

We accessed published whole-gene resequencing data from 87 individuals of *J. regia* from Ding *et al*. (2022); Ji *et al*. (2021), 60 individuals of *J. mandshurica* from (Xu *et al*. 2021), and 26 *J. regia*, 11 *J. hindsii*, 12 *J. microcarpa*, and 13 *J. nigra* from Stevens *et al*. (2018). Published resequencing data for *Cyclocarya* and *Platycarya* were obtained from Qu *et al*. (2023) and Zhang *et al*. (2019), respectively. We used publicly available long-read genome assemblies of plants within *Fagales* generated by Zhu *et al*. (2019); Marrano *et al*. (2020); Fitz-Gibbon *et al*. (2023); Zhang *et al*. (2021); Lovell *et al*. (2021); Zhang *et al*. (2023); Ding *et al*. (2023); Ning *et al*. (2020); Li *et al*. (2022a); Zhang *et al*. (2020); Sork *et al*. (2022); Qu *et al*. (2023); Cao *et al*. (2023); Guzman-Torres *et al*. (2023).

We accessed mRNA transcriptome sequence data from floral tissues of *J. mandshurica* from Qin *et al*. (2021) and Li *et al*. (2022b). We accessed small RNA sequencing libraries from floral buds of a protogynous and protandrous individual of *J. mandschurica* from Li *et al*. (2023).

We accessed whole-genome resequence data for 34 varieties of *C. illinoinensis* from Xiao *et al*. (2021). We verified the variety identity for 12 of these samples by comparison of genetic relatedness at overlapping sets of SNPs to reduced representation sequence data from 83 varieties of cultivated pecan from Bentley *et al*. (2019). The remainder of these samples were not used in the analysis, as several were labelled as varieties that did not match their predicted identity using data from Bentley *et al*. (2019). We chose the data from Bentley *et al*. (2019) as the standard as dichogamy type was rigorously documented for the trees in this data set. Whole-genome resequencing for 15 *Carya* individuals of different species were obtained from Huang *et al*. (2019).

### Sequence alignment, variant calling, and coverage analyses

We mapped short read sequencing libraries to available long-read reference genomes using *bwa* 0.7.17 (Li and Durbin 2009) with default parameters. To examine structural differences between G-locus haplotyes, we used BLAST (Altschul *et al*. 1990) to identify and align syntenic regions containing *TPPD-1* orthologs. We used both minimap2 (Li 2018) and Anchorwave (Song *et al*. 2022) to align entire chromosomes and G-locus regions for variant calling and divergence calculations. In one analysis, we aligned all available genomes to the Walnut 2.0 (*g* haplotype) (Fig. S7). For divergence calculations, we aligned genomes containing alternate G-locus haplotypes within species for *J. regia*, *J. californica*, and *J. mandschurica* using Anchorwave. As Anchorwave uses a genome annotation in the alignment, we used *liftoff* (Shumate and Salzberg 2021) to port the Walnut 2.0 annotation onto assemblies lacking an annotation. Variant calling was done with *bcftools* 1.17 (Danecek *et al*. 2021). Variants were filtered in *vcftools* 0.1.16 (Danecek *et al*. 2011). Filters were set at minDP 10 and minGQ 30 for most analyses, but were adjusted on a per-analysis basis. We measured read depth in windows across G-loci in *Juglans* and *Carya* using *samtools depth* (Danecek *et al*. 2021). Read depth was normalized for each individual by the average read depth across a different chromosome which was chosen arbitrarily.

We used Salmon (Patro *et al*. 2017) to align RNA-seq data transcripts to a transcriptome of *J. mandshurica* and *tximport* (Soneson *et al*. 2015) to quantify normalized transcript abundances. We used STAR 2.7.6 (Dobin *et al*. 2013) to align transcripts to the reference genome to measure relative allelic depths at *J. mandshurica* G-locus SNP positions with fixed differences. SNPs were ascertained from whole-genome resequencing of 60 *J. mandshurica*, where we phased variants using Beagle 5.4 (Browning *et al*. 2021) with a dummy SNP at the location of the *GJ1* indel. For small RNA sequence data, we first trimmed adapter sequences using *skewer* (Jiang *et al*. 2014) and filtered for reads 18-36 bp in length. Reads were aligned to an assembly of the protogynous assembly of *J. mandshurica* using *bowtie* 1.3.0 (Langmead *et al*. 2009). We used two mapping approaches, one which reported only uniquely-mapping reads with one mismatch (-v 1 –m 1), and one which reported reads that map to at most 10 locations in the genome with a single mismatch (-v 1 –m 10).

### GWAS and linkage disequilibrium

We performed GWAS for dichogamy type separately in 44 individuals of *J. hindsii*, 26 *J. regia* individuals, and 30 *Carya illinoinensis* individuals. GWAS was done in GEMMA 0.98.3 (Zhou and Stephens 2012), which controls for genome-wide relatedness among samples. We calculated genotypic LD as the *r*^2^ value between genotypes at pairs of loci using *vcftools*, filtering sites for a minimum distance of 100 bp or 5 kb.

### Phylogenetic analyses and synteny

To construct a phylogeny of *GJ1* repeats and their homologs, we extracted the genomic coordinates of individual repeat subunits from pairwise BLAST alignments. We aligned subunits a, b, and c separately using *muscle* (Edgar 2004) and then concatenated alignments. We constructed a maximum-likelihood phylogeny using *IQ-Tree* (Nguyen *et al*. 2015) and obtained node support values using the program’s ultrafast bootstrap approximation algorithm. We constructed a species phylogeny for *Carya* genome assemblies and *J. regia*, from concatenated alignments of 12,101 single copy orthologs identified in OrthoFinder (Emms and Kelly 2019). The phylogeny of the *Carya* G-locus was inferred similarly using just the 20 genes within the G-locus, as well as for each gene individually. We inferred and visualized synteny across the *Carya* G-locus using MCscan (Tang *et al*. 2008) which leverages collinearity of orthologs.

While G-locus haplotypes show strong conservation of synteny, we note a small number of predicted genes that are not shared between haplotypes (see Fig. 3F). None of these was annotated independently in two assemblies. However, we investigated whether homologous sequence in other assemblies may have been missed by the annotation pipeline. We therefore used BLAST to search for sequence homologous to these uniquely annotated genes in other *Carya* G-locus assemblies (Table S5). For two genes that were uniquely annotated in the *C. sinensis* assembly, we found significant BLAST hits within syntenic regions of the ‘Lakota’ primary assembly (Fig. 3F), but not within G-locus regions of the other assemblies (although we find hits elsewhere in the assemblies). This result suggests that these are ancient duplications, perhaps with conserved function. However, further validation of these predicted genes with gene expression and additional de novo assemblies is needed to fully address the role of gene duplications in the assembly of G-locus haplotypes.

### Haplotype nucleotide divergence and dating

We calculated nucleotide divergence across the region encompassing the *Juglans* G-locus in 500 bp windows from genome alignments using custom R scripts. We find that the divergence between G-locus haplotypes is comparable to the divergence between *Juglans* and *Carya*, indicating deep divergence of *Juglans* G-locus haplotypes, potentially occurring close in time to the split between *Juglans* and *Carya* (*∼*70 Myr). We used a molecular clock approach with substitution rates estimated from nonsynonymous coding regions in *Juglans* (Ding *et al*. 2023; Zhu *et al*. 2019). Here, we used alignments between G-locus haplotypes within three species (*J. regia*, *J. californica*, *J. mandschurica*), and took the average of the maximum divergence value within any 500 bp window on either side of the G-locus indels within 4 kb. We then adjusted this average value of *D_XY_* for multiple hits using the Jukes and Cantor (1969) distance correction. We then calculated the divergence time as *T* = *D_XY_ /*(2*µ*). Using two reported estimates of the substitution rate (*µ* = 1.5 *×* 10*^−^*^9^ per bp per year (Ding *et al*. 2023), and 2.5 *×* 10*^−^*^9^ per bp per year (Zhu *et al*. 2019)), we obtain estimates of 68.8 Mya and 41.3 Mya, respectively. We note there is considerable uncertainty about the substitution rate for this region. Furthermore, evidence of rare recombination between haplotypes within the *TPPD-1* sequence over deep timescales (Fig. S5) suggests that the genealogies of these linked regions in contemporary G-locus haplotypes may differ from the genealogy of the causal variants. Nonetheless, these estimates accord well with the fact that the divergence between *Juglans* G-locus haplotypes is comparable in magnitude to the divergence between *Juglans* and *Carya* in this region (Fig. S6). We also note that it is consistent with trans-specific SNPs across the deepest split within *Juglans*, the inferred *GJ1* phylogeny, and read depth analyses in supporting haplotype divergence in the common ancestor of *Juglans*.

To calculate divergence between pecan G-locus haplotypes, we aligned coding regions from the ‘Lakota’ and ‘Pawnee’ primary assemblies and calculated *D_XY_*using *pixy*. *pixy* was also used to calculate individual-level heterozygosity in other *Carya* species at pecan G-locus SNPs and SNPs that differentiated the two East Asian *Carya* G-locus haplotypes, and to calculate heterozygosity for *Pterocarya* and *Cyclocarya* individuals across regions syntenic with *Juglans* and *Carya* G-loci. We used the R package *ape* (Paradis and Schliep 2019) to calculate divergence at fourfold degenerate sites within aligned coding sequences from the *Carya* G-locus. To estimate a divergence time, we used the ratio of the *D_XY_* between *Carya* haplotypes and from the average of both *Carya* haplotypes to *J. regia*, adjusting divergence values for multiple hits using Jukes and Cantor (1969).

### Polymorphism within haplotypes and tests of selection

We examined polymorphism within Glocus haplotypes in a sample of 113 *Juglans regia*, containing individuals sampled from across the species’ range and without reference to dichogamy type (Ding *et al*. 2022; Ji *et al*. 2021; Stevens *et al*. 2018). In this sample, we observed 60 *gg*, 50 *Gg*, and 2 *GG* individuals for the G-locus structural variant. Thus, proportions of protandrous and protogynous individuals are close to 50/50 in this broad sampling (P=0.46 under null hypothesis of equal proportions). The proportion of *GG* genotypes is lower than expected under Hardy-Weinberg equilibrium (P=0.036), consistent with disassortative mating at the G-locus or selection against *GG* genotypes.

In *Juglans*, to test for an excess of nonsynonymous fixed differences between G-locus haplotypes relative to polymorphism (or conversely, an excess of nonsynonymous polymorphism), we performed a McDonald-Kreitman test for *TPPD-1* coding sequence (McDonald and Kreitman 1991) in sample of 113 *J. regia* and in a sample of 46 *J. hindsii*. To obtain data partitioned by haplotype, variants were first filtered for a minor allele count of 2 and then phased haplotypes using Beagle 5.4 (Browning *et al*. 2021). In order to assign haplotype identities to phased haplotypes, we added a line to the VCF with a dummy SNP representing an individual’s genotype for the G-locus structural variant. In *J. regia*, we discarded one heterozygote which showed an erroneous phase switch between its two haplotypes; aside from this we saw no other evidence of phasing errors *TPPD-1* in heterozygotes in either species sample. Furthermore, phasing resulted in two distinct clusters of haplotypes with non-overlapping distributions of the number of variants compared to the reference. We used a custom R script to calculate nonsynonymous and synonymous polymorphism and divergence in the sample, and performed the MK test using Fisher’s exact test. We ignored singletons within haplotype groups in determining fixed differences between haplotype groups.

In *J. regia*, we found 9 fixed nonsynonymous variants and 6 fixed synonymous variants in *TPPD-1* coding sequence. Using a long read alignment between *J. regia* and an outgroup (pecan) to polarize SNPs, we identified 5 nonsynonymous and 3 synonymous fixed differences derived in the *J. regia* G lineage, 4 nonsynonymous and 2 nonsynonymous derived in the *g* lineage, and one synonymous site with an ambiguous ancestral state. We found limited polymorphism within haplotype groups in *TPPD-1* coding sequence: 2 nonysnonymous and 1 synonymous in *g* haplotypes, and 4 nonysnonymous and 1 synonymous in *G* haplotypes. In *J. hindsii*, we observed 8 nonsynonymous and 5 synonymous fixed differences between haplotypes. We observed a near complete lack of polymorphism within *TPPD-1* coding sequence in this population, with only one polymorphism segregating in multiple *g* haplotype copies, and zero polymorphisms segregating in multiple *G* haplotype copies.

In pecan, we similarly phased SNPs in coding regions across the *Carya* G-locus along with with a dummy SNP placed in the center of the G-locus to represent the structural variant. We saw no evidence of phasing errors by checking for haplotype switching in heterozygotes. Using *J. regia* as an outgroup to polarize SNPs, we identified 235 G-locus coding SNPs that fixed in G lineage, 118 of these nonsynonymous changes. We identified 215 that fixed in g lineage, 103 of these nonsynonymous changes. Fifteen remaining fixed differences had an ambiguous ancestral state where *J. regia* showed a different allele than either pecan haplotype. We tested for a difference in 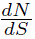 values between the *G* and *g* lineages using a Chi Squared test, which did not yield a statistically significant result.

We used *pixy* to estimate pairwise nucleotide divergence (*π*) between the sets of homozygous individuals within coding sequence at the *Carya* G-locus (*gg*: 13, *GG*: 2). The two *GG* indivduals are not known to be close relatives from pecan pedigree records, and they do not show an unusually high kinship coefficient compared to other pairs of individuals in our analysis set. To test for significance, we estimated *π* for all combinations of two *gg* individuals, and checked whether in any case the estimated value was equal to or lower than the value observed for *GG* individuals. We separately estimated Watterson’s Theta from phased haplotypes and found an 8.7-fold reduction for *G* haplotypes compared to *g* haplotypes (Θ*_W,g_*= 0.00106, Θ*_W,G_* = 0.00012).

### PacBio IsoSeq

Tissues were collected from various locations at the University of California in Davis and the USDA’s National Clonal Germplasm Repository in Winters, CA. Collected tissue was wrapped in foil and immediately immersed in liquid nitrogen in the field to preserve RNA quality. Frozen tissues were subsequently pulverized in liquid nitrogen in a mortar and pestle for extraction.

The extraction buffer used was 4M guanidine isothiocyanate, 0.2 M sodium acetate pH 5.0, 2mM EDTA, 2.5% (w/v) PVP-40. To this we added 400 uL Lysis buffer to 100 mg tissue and homogenize with pestle for 30 seconds, then 600 uL Lysis buffer (1 mL total)was added and vortexed for 20 seconds. From this, 500 uL of the homogenate was processed using the RNeasy Plant Mini kit (Qiagen) according to manufacturer’s protocol (centrifugations steps were performed at 12,000 rpm for 30 seconds). RNA was eluted with 50 uL nuclease-free water at Step 9 in manufacturer’s protocol. This was followed by DNase digestions with Turbo DNA-free kit (Invitrogen) according to manufacturer’s protocols. Finally, RNA samples were cleaned up using HighPrep RNA Elite beads (MagBio Genomics) according to the manufacturer’s 96 well format protocol for 10 uL reaction volume. Final elution was performed with 20 uL of nuclease-free water that was heated at 60C for *∼*10 minutes before adding to sample.

For quality control and quantitation, samples were subsequently checked for purity on Nanodrop, quantified on Qubit, and checked on Bioanalyzer.

SMRTbell libraries were constructed and sequenced at the University of California Davis Genome center. Sequencing was performed on the Pacbio Sequel II. Demultiplexing and post processing of sequence data to create high quality full length non-chimera consensus transcripts (FLNC) was performed using the PacBio bioinformatics pipeline (ccs v4.2.0, lima v1.11.0, isoseq v3).

To determine the presence of a transcribed sequence in an IsoSeq library of (full-length non-chimeric) FLNC reads, command line blastn 2.12.0+ was used with default parameters. The sequence for the *TPPD-1* was queried against each individual library separately. Matches with a percent identity greater than or equal to 90 were considered positives.

### Transposable Element Annotation

We performed de-novo whole genome annotation of transposable elements (TEs) for long-read assemblies of both G-locus haplotypes in *J. regia*, *J. californica*, and *J. mandshurica* using EDTA (Ou *et al*. 2019), and compared coordinates of annotated TEs to the coordinates of the derived *gJ3* insertion. EDTA predicted the presence of a CACTA-like DNA transposon at the coordinates of *gJ3*. We separately identified the presence of terminal inverted repeats near the *gJ3* insertion endpoints and evidence of target site duplication.

We also used EDTA to annotate assemblies of ‘Pawnee’ and ‘Lakota’ pecan as well as assemblies of *Carya sinensis* and *C. cathayensis*. We computed the proportion of sequence covered by predicted TEs for three categories – (1) within the G-locus (defined by endpoints of genes falling inside the pecan GWAS peak, (2) in 300kb of sequence surrounding the G-locus (150kb on either side), and (3) across the whole genome.

## Supplementary Material

**Figure S1:**
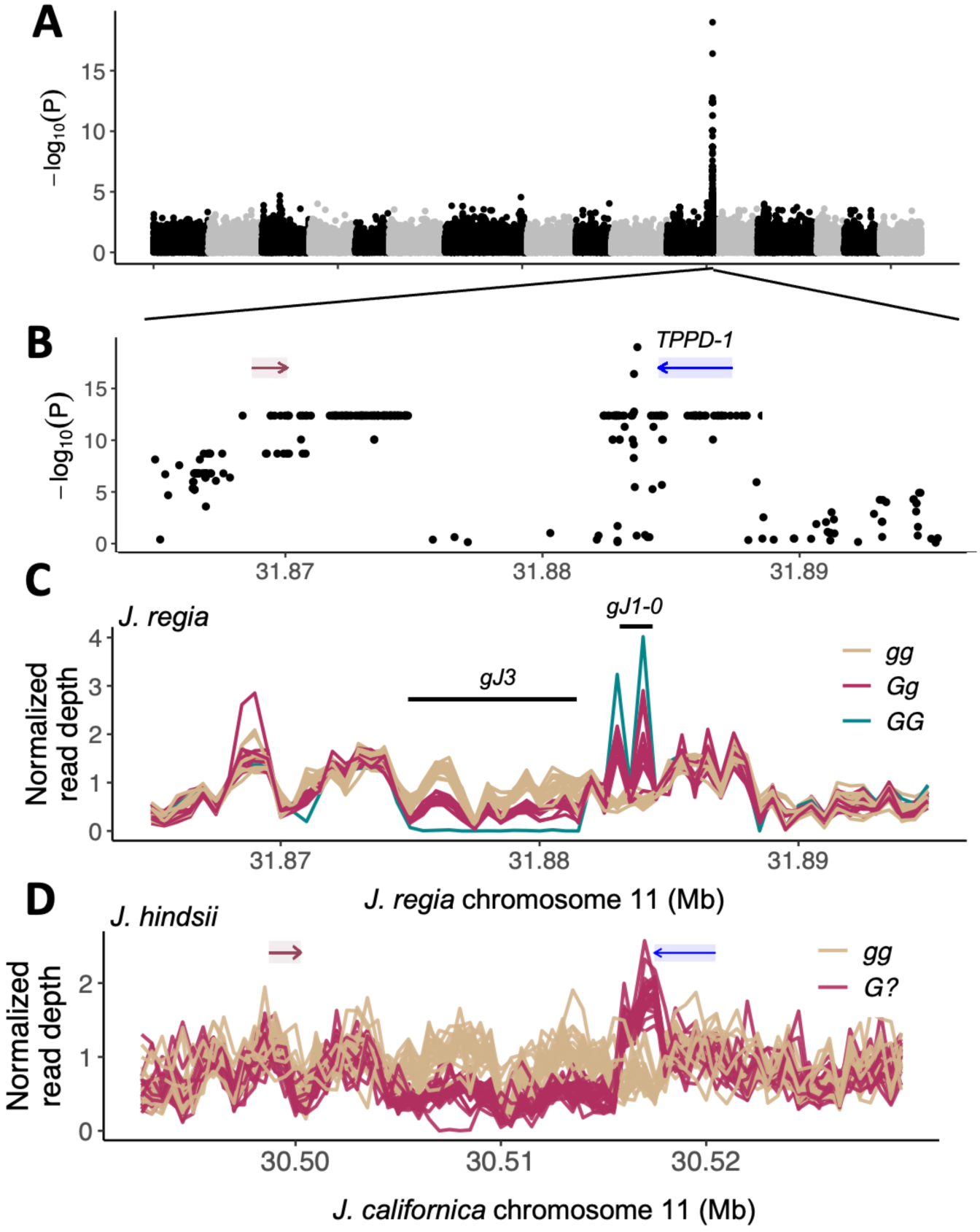
**A**) GWAS for dichogamy type in 26 *J. regia* individuals. **B)** Zoomed in region of strong association. Here, read mapping and variant calling was done using the Walnut 2.0 genome, an assembly from the ‘Chandler’ variety, a protandrous tree (*gg* genotype.) Highly similar results were seen mapping to an independent assembly of a protandrous variety (‘Serr’, Zhu *et al*. 2019) **C)** Normalized average read depth in 1 kb windows for 26 *J. regia*individuals of known genotype across the Walnut 2.0 assembly (*g* haplotype). Black bars indicate the position of the (*gJ3*) indel and *gJ1-0* (See Fig. 2 in main text for notation of structural variants). The spike in coverage at *gJ1-0* seen for protogynous individuals corresponds to sequencing reads originating from repeats within the *GJ1* insertion in the *G* haplotype.

**Figure S2:**
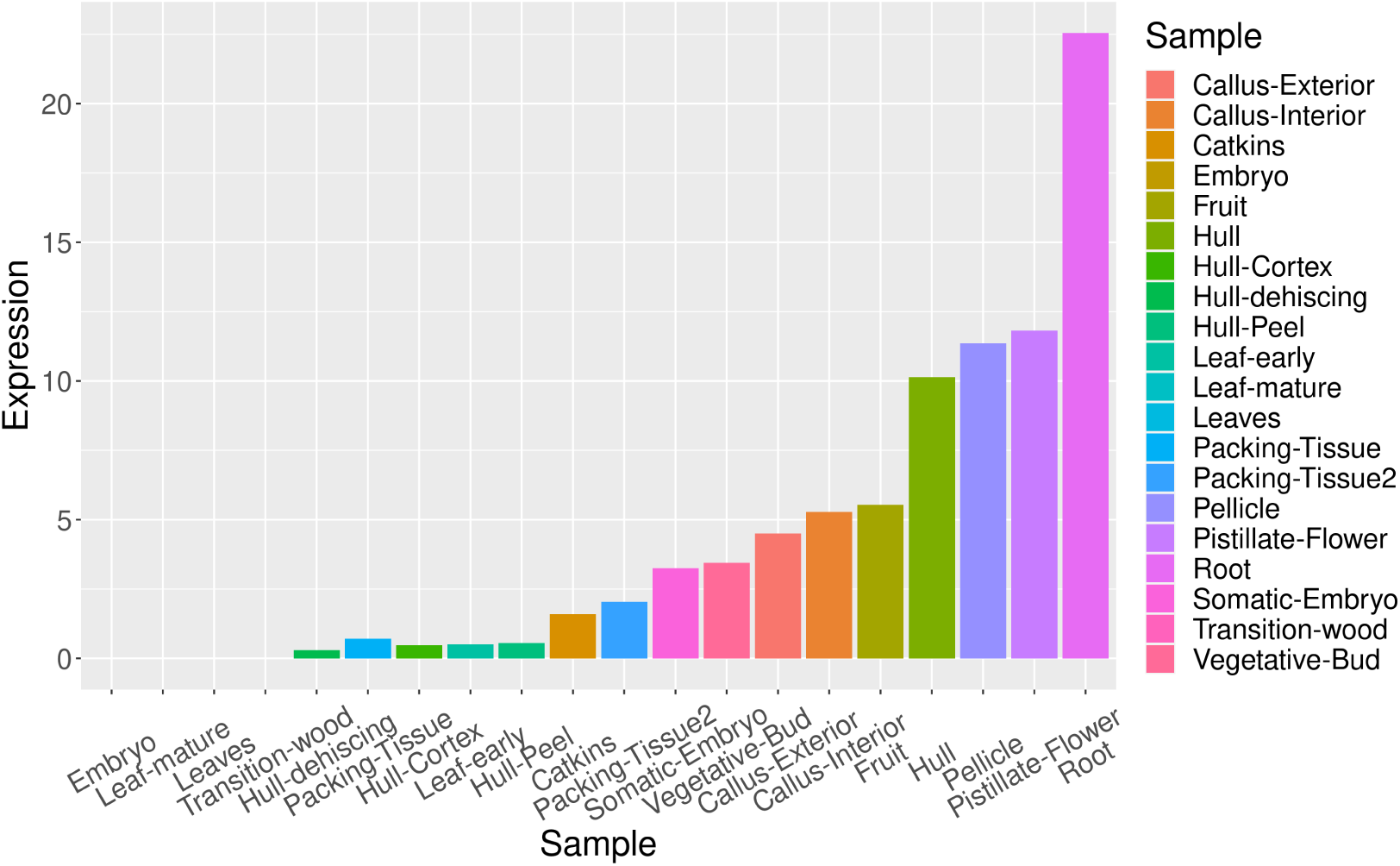
Gene expression levels of *TPPD-1* (LOC108984907) from published short read RNA-seq libraries of *J. regia*‘Chandler’ (*gg* genotype). Data from (Chakraborty *et al*. 2016). Y-axis measures Fragments Per Kilobase of transcript per Million mapped reads.

**Figure S3:**
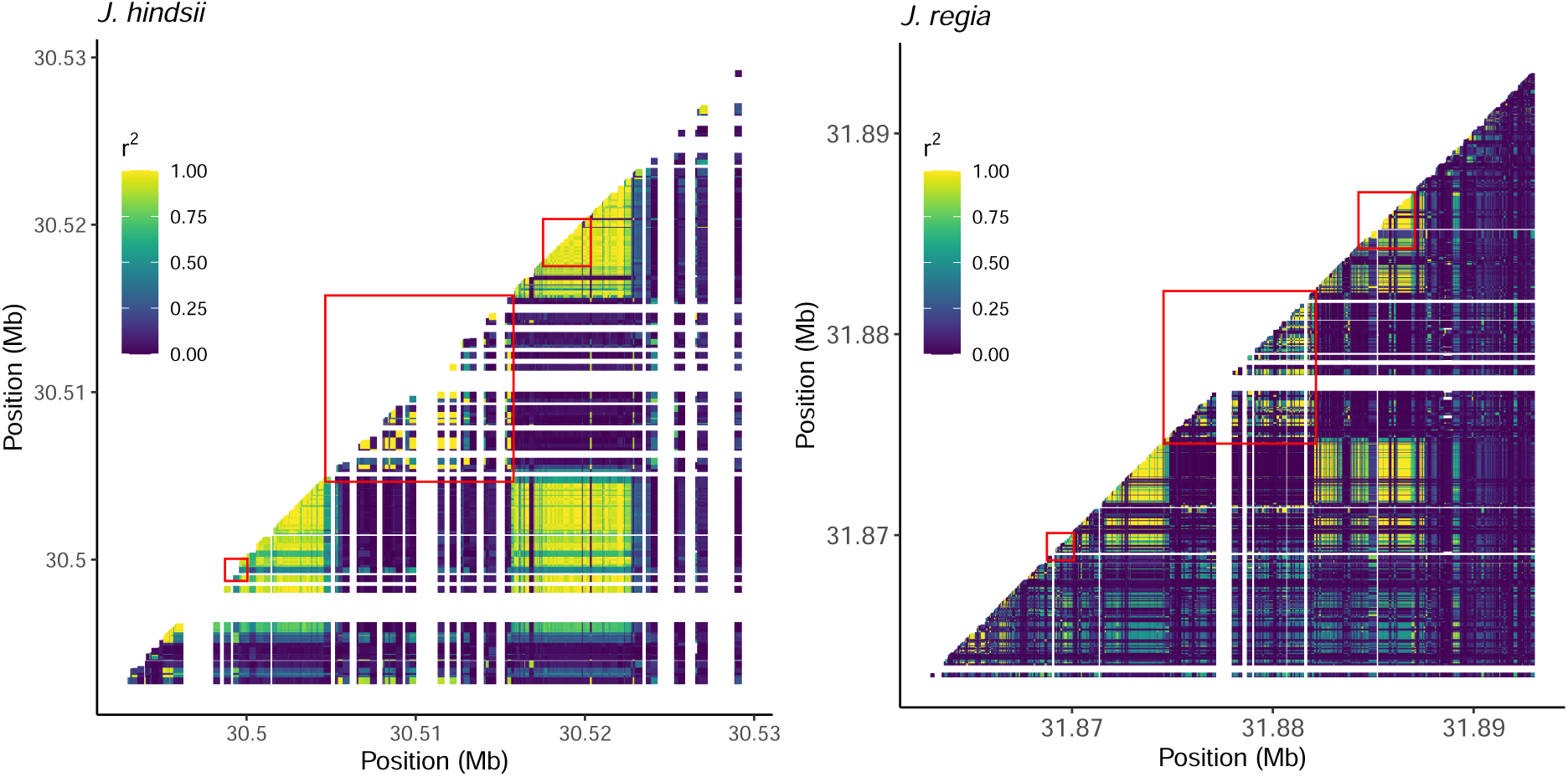
Correlation of diploid genotypes between loci (LD) in the region surrounding the G-locus in two species of *Juglans*. **Left)** LD in a sample of 46 from a natural population of *J. hindsii*. Shown for reads mapped to an assembly of the sister species *J. californica* which was identified as the *g* haplotype (same assembly as in Fig. S1D, but not Fig. 1E). For this analysis we filtered variants for a minimum minor allele freuqency of 0.1. **Right)** LD in a sample of 113 *J. regia* (see methods for data sources). Shown for reads mapped to the Walnut 2.0 assembly (‘Chandler’, *gg*) Red boxes from bottom left to top right indicate positions of a *NDR1/HIN1* –like gene, the *gJ3* indel, and *TPPD-1*. See Fig. 2A,B for schematic of indels. Note that sequence from *G* haplotypes is absent within *gJ3*, and only *g* haplotypes contribute to the signal of LD for a pair of sites where one site falls within *gJ3*. The low levels of LD for pairs of sites where one site is within *gJ3* indicates that recombination is not reduced across this region between *g* haplotypes relative to the genomic background.

**Figure S4:**
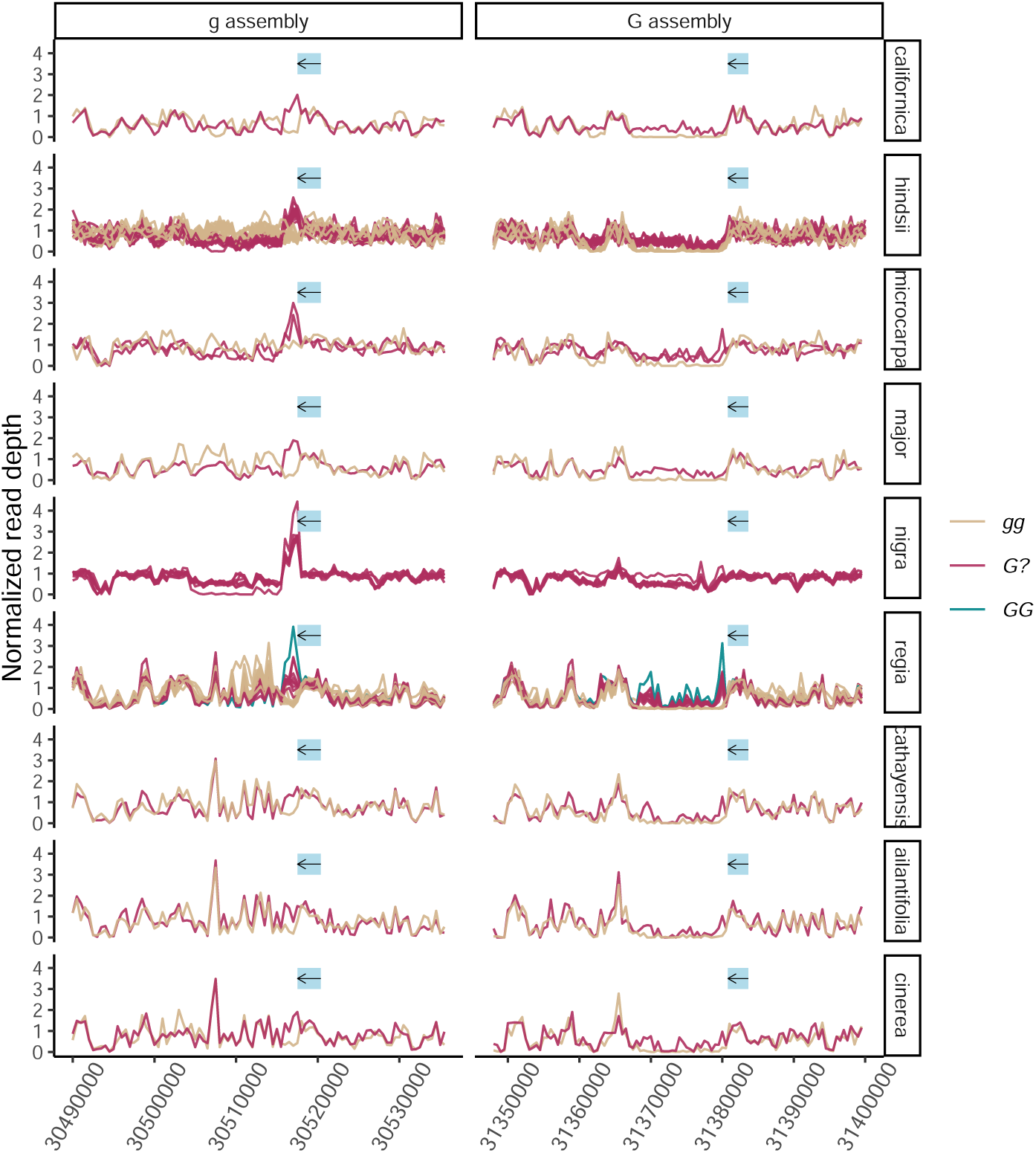
Patterns of read depth from multiple species against two pseudohaploid assemblies of *J. californica* at the region syntenic with the GWAS hit in *J. regia* reveal the same segregating structural variant across multiple species in association with heterodichogamy flowering type. The gray bar indicates the location of the TPP ortholog in each assembly, and the arrow indicates direction of transcription. Read depth was normalized by the average coverage across an arbitrarily chosen separate chromosome. Protandrous individuals are known genotype *hh*, while the genotype of protogynous individuals is not known a priori without pedigree information. One homozygote for the protogynous allele is known in *J. regia* (variety ‘Sharkey’, Gleeson (1982)). Read depth for the *J. nigra* ‘Hay’ variety also indicate it is homozygous for the protogynous allele.

**Figure S5:**
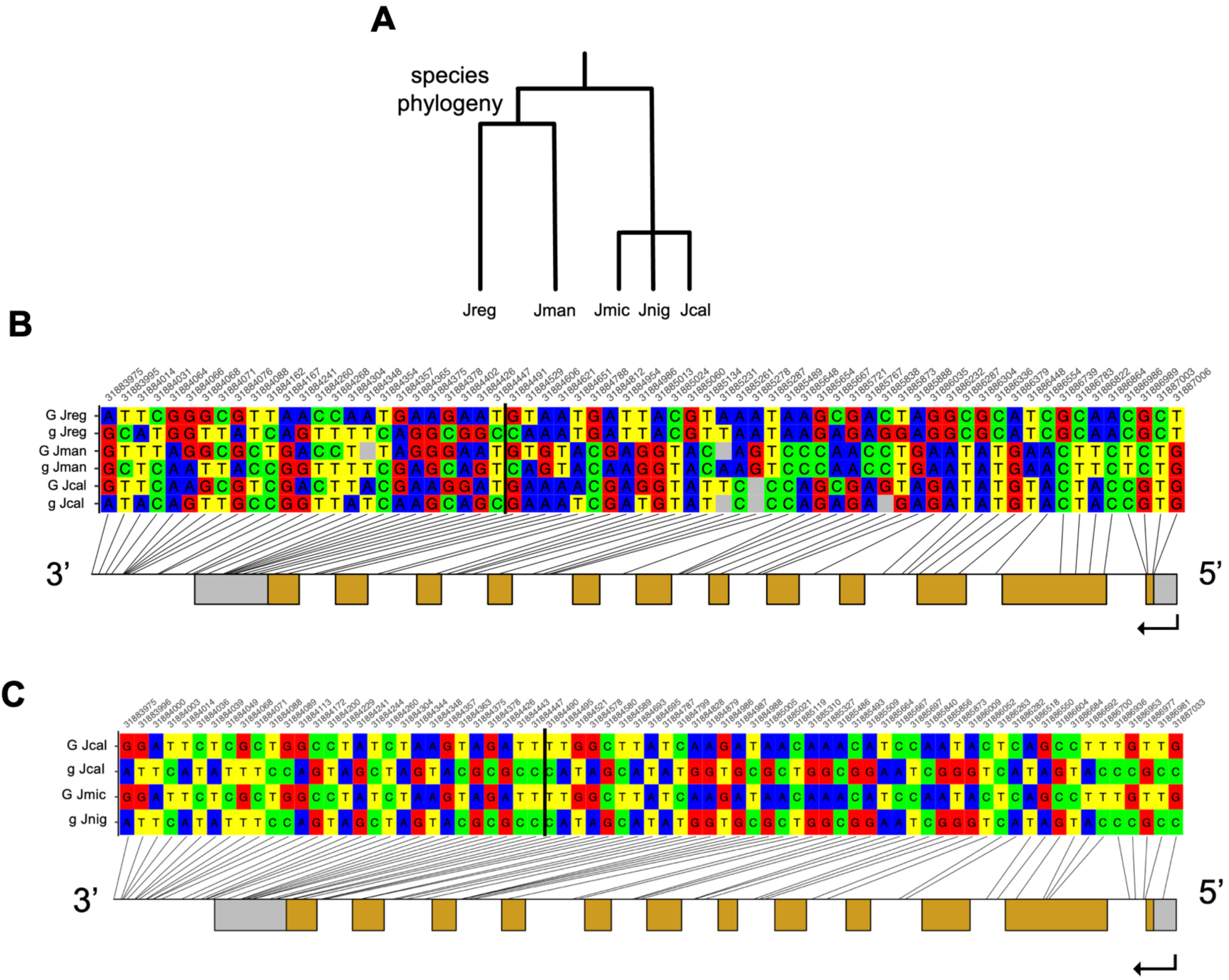
Polymorphism within the *TPPD-1* gene and its 3’ region (*GJ1-0*) in *Juglans* G-locus haplotypes. **A)** Species phylogeny for *J. regia*, *J. mandschurica*, *J. microcarpa*, *J. nigra*, and *J. californica*, adapted from Mu *et al*. (2020); Zhou *et al*. (2021). **B)** Polymorphism in G-locus haplotypes for 3 species reprenting the two deepest splits in the *Juglans* phylogeny (i.e. *∼*35-50 Myr ago). Numbers along top indicate positions of nucleotides in the Walnut 2.0 reference genome. Singletons are omitted from the sample. Diagonal lines connect SNP genotypes to their locations within the gene model of *TPPD-1*. Below, rectangles indicate exons – gray rectangles represent UTRs, and brown rectangles represent CDS. SNPs that consistently group by haplotype in this sample are localized to the 3’ UTR or within *GJ1-0*, but within *TPPD-1* CDS, polymorphisms quickly transition to grouping by species. The black vertical line indicates the border between the 3’UTR and the CDS of the last exon in the reference annotation. **C)** As in (B), but for G-locus haplotypes in three species of North American black walnuts (sect. *Rhysocaryon*, *∼*10 Myr divergence). The phylogenetic relationships between these taxa are not well-resolved. In this case, trans-specific polymorphism extends across the *TPPD-1* coding region.

**Figure S6:**
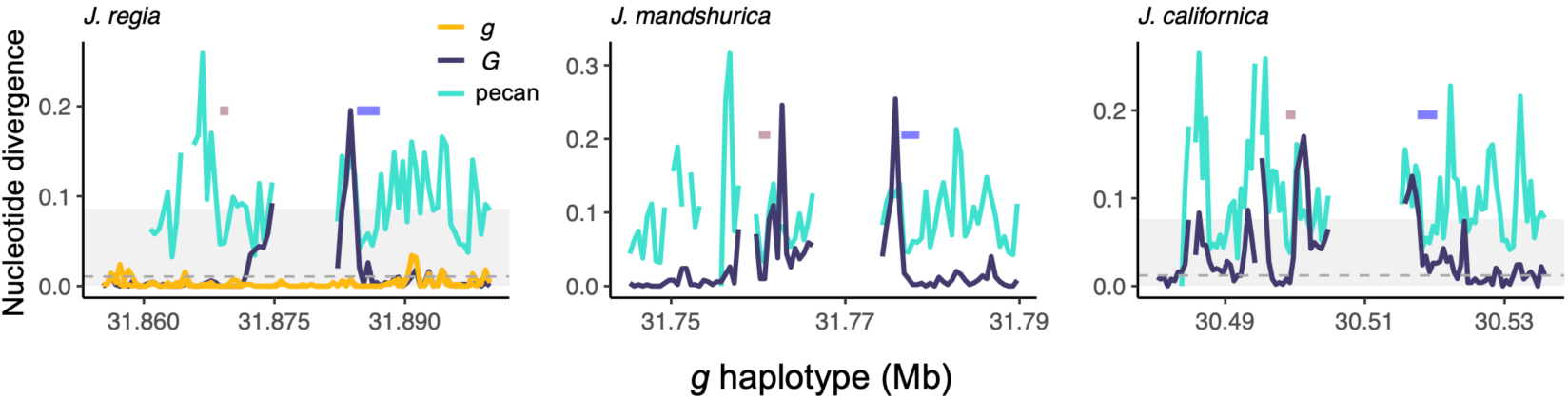
Nucleotide divergence of *Juglans* G-locus haplotypes in comparison with nucleotide divergence between *Juglans* and *Carya*. Nucleotide divergence is shown for 500 bp windows calculated from genome alignments of both G-locus haplotypes in three *Juglans* species spanning the two deepest divergence events in the genus. Nucleotide divergence between alternate haplotypes is comparable that observed between *Juglans* and *Carya*.

**Figure S7:**
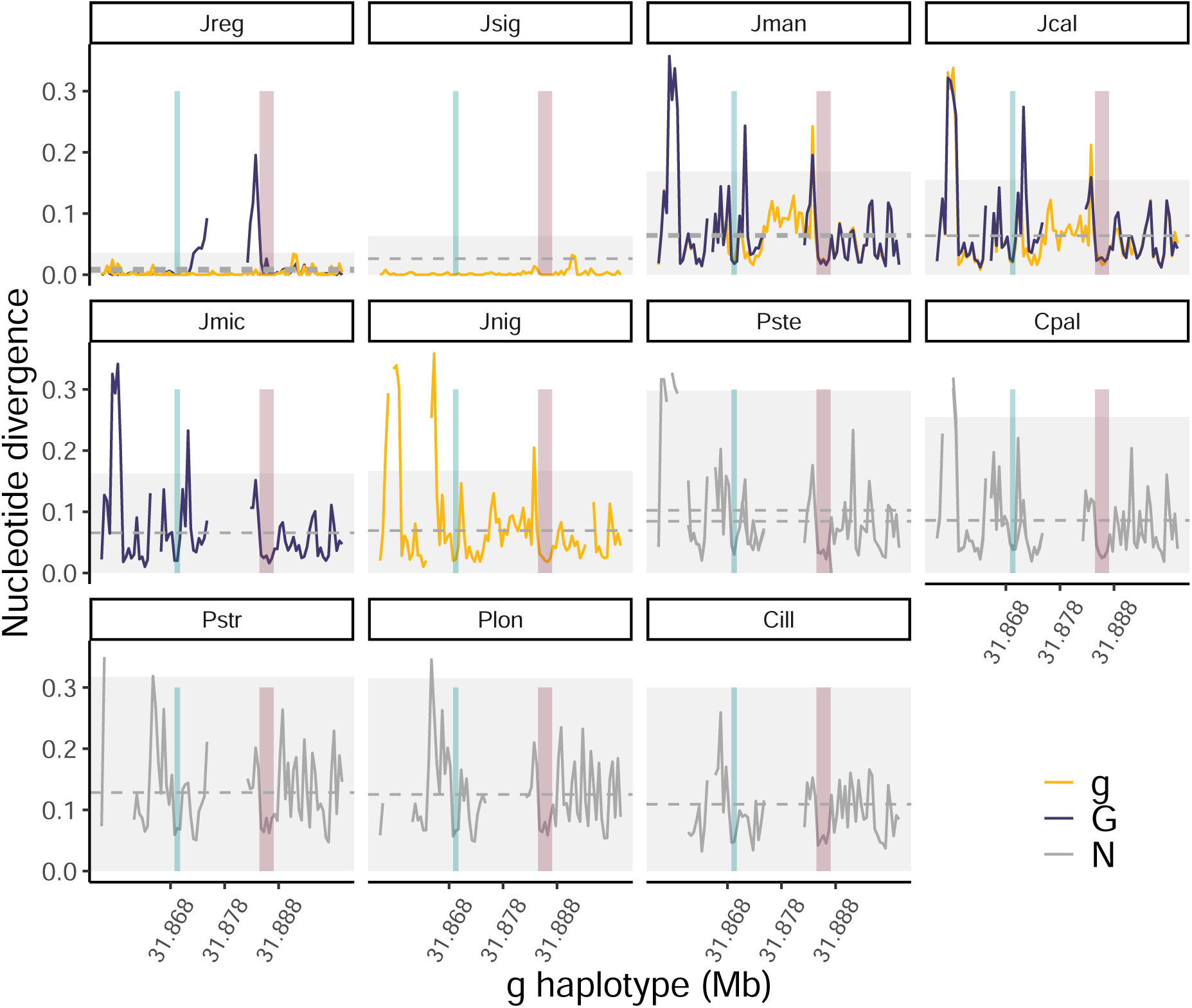
Nucleotide divergence of Juglandaceae genomes across the *g* haplotype of *J. regia* ‘Chandler’, calculated in 500bp windows from whole-genome alignments. Dotted gray lines show the average calculated over the entire aligned chromosome. Shaded gray interval indicates the top 95% quantile.

**Figure S8:**
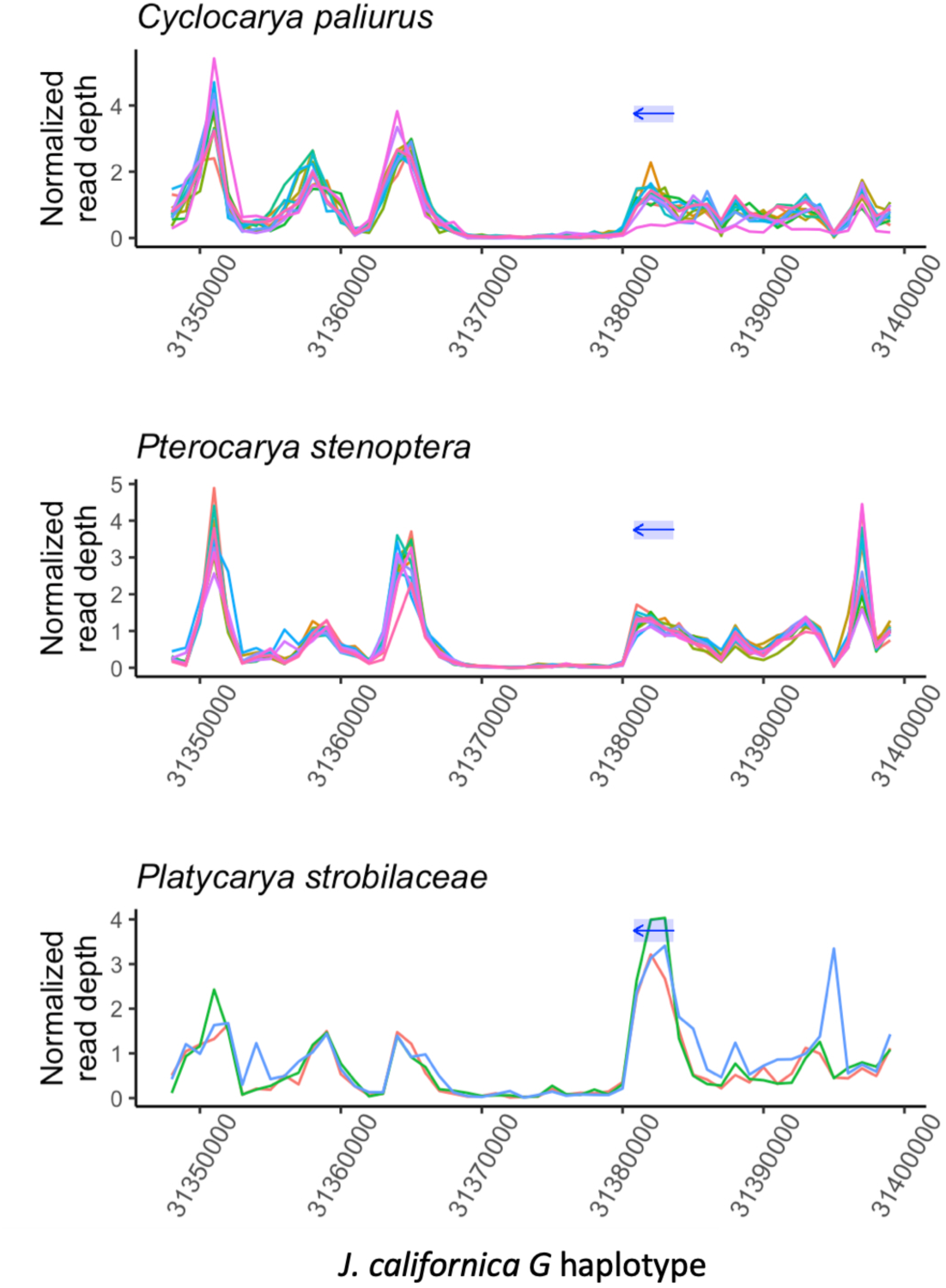
Normalized average read depth in 1kb windows for 12 individuals of *Cyclocarya paliurus* (sample contains both protandrous and protogynous individuals, Qu *et al*. 2023), 13 *Pterocarya stenoptera*, and 3 *Platycarya strobilaceae* against an assembly of the *Juglans G* haplotype from *J. californica*. Blue rectangle and arrow indicates the position of *TPPD-1* in the assembly.

**Figure S9:**
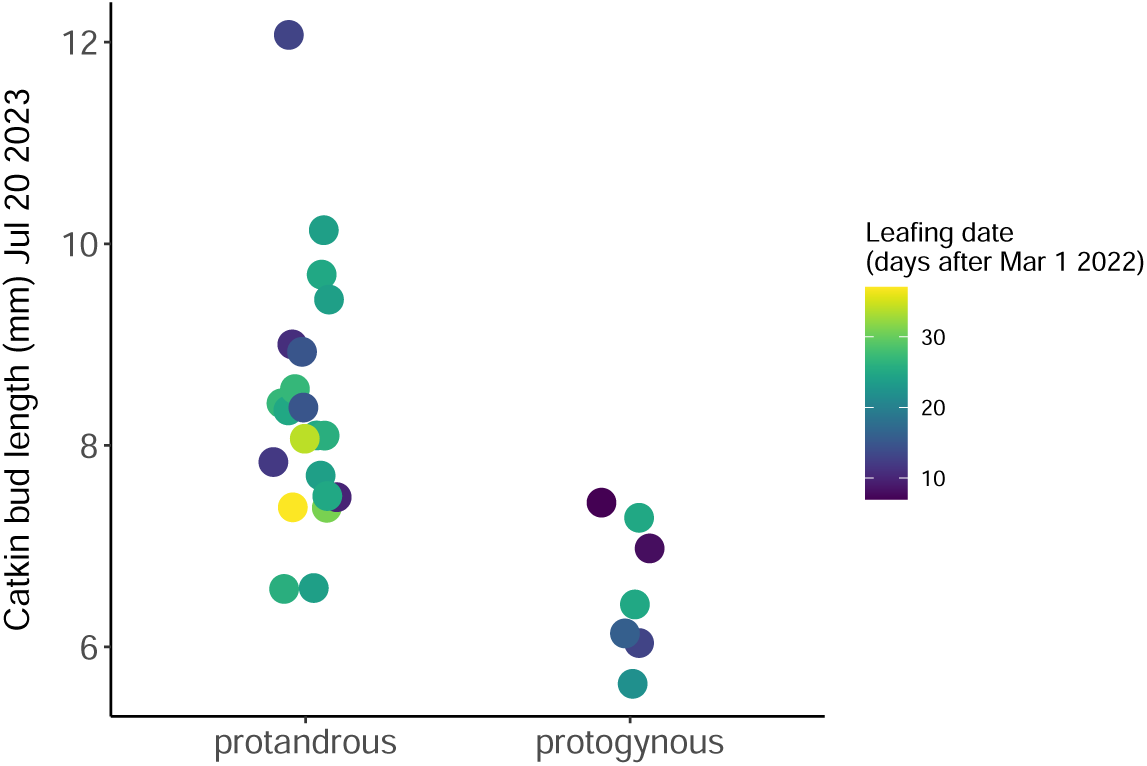
Protandrous and protogynous morphs of *J. regia* differ in size of male catkin buds in the season prior to flowering (*β* = 2 mm, P*<*0.001). We fit a linear model predicting catkin bud length from dichogamy type with leafing date as a covariate. While leafing date was not a significant predictor, we note that the slope estimate is in the expected direction. Notably, the earliest leafing protogynous varieties have catkin buds roughly of similar size to the latest leafing protandrous varieties.

**Figure S10:**
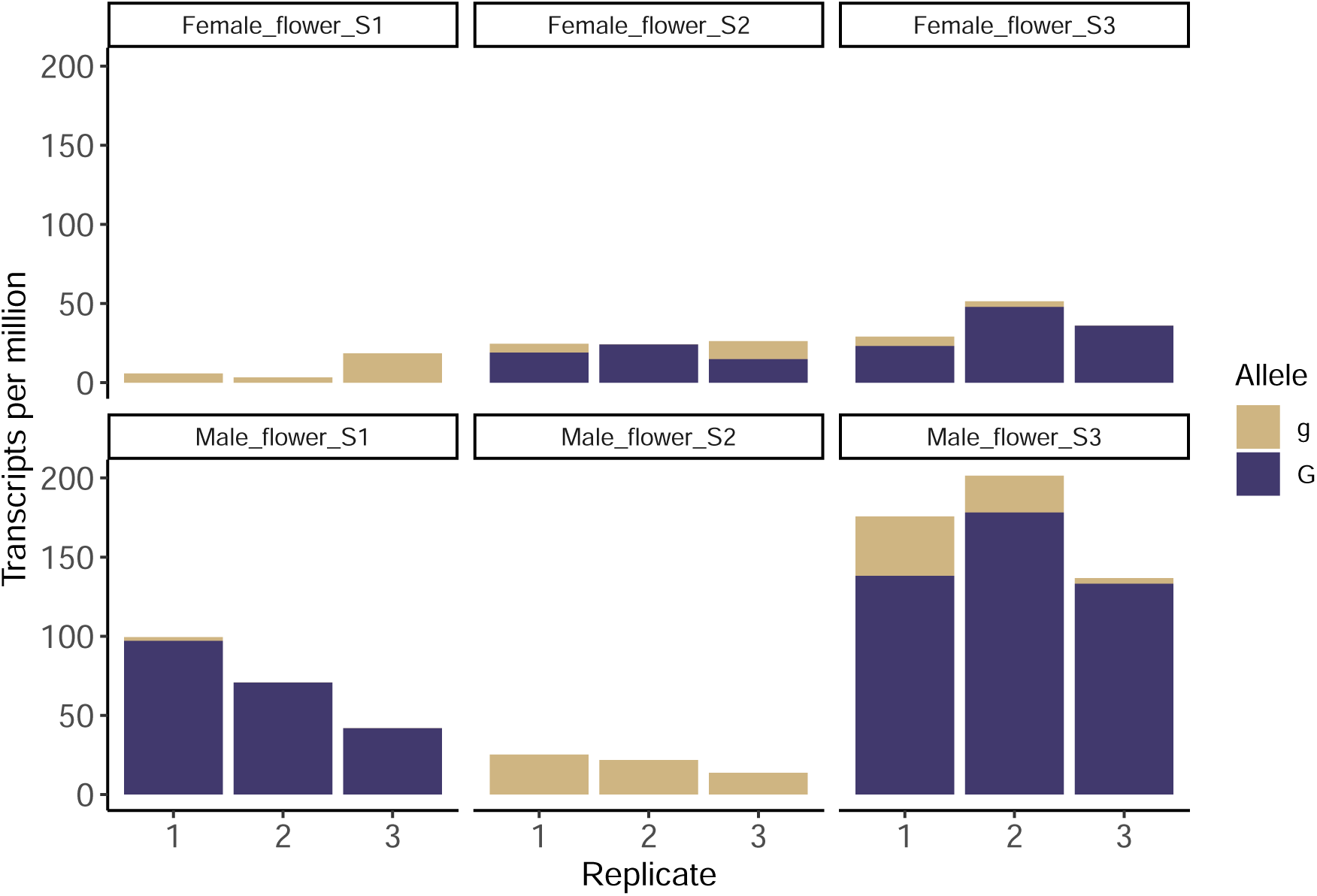
Reanalysis of RNA-seq experiment from Li *et al*. (2022b). The authors generated RNA-seq from male and female inflorescences at three developmental stages: S1 – dormant buds collected in the season prior to flowering; S2 – season of flowering, prior to anthesis; S3 – mature flowers. We measured relative read depth in each sample at SNPs within *TPPD-1* that are fixed between *J. mandshurica* G-locus haplotypes, ascertained from whole-genome resequencing data 31 individuals of this species (Zhang *et al*. 2021). We verified that the difference in allele-specific depth is not a mapping artefact by mapping the reads to assemblies of both haplotypes. Each unique tissue is described to have 3 biological replicates, although relatedness estimates indicate that a subset of samples may represent the same genotype.

**Figure S11:**
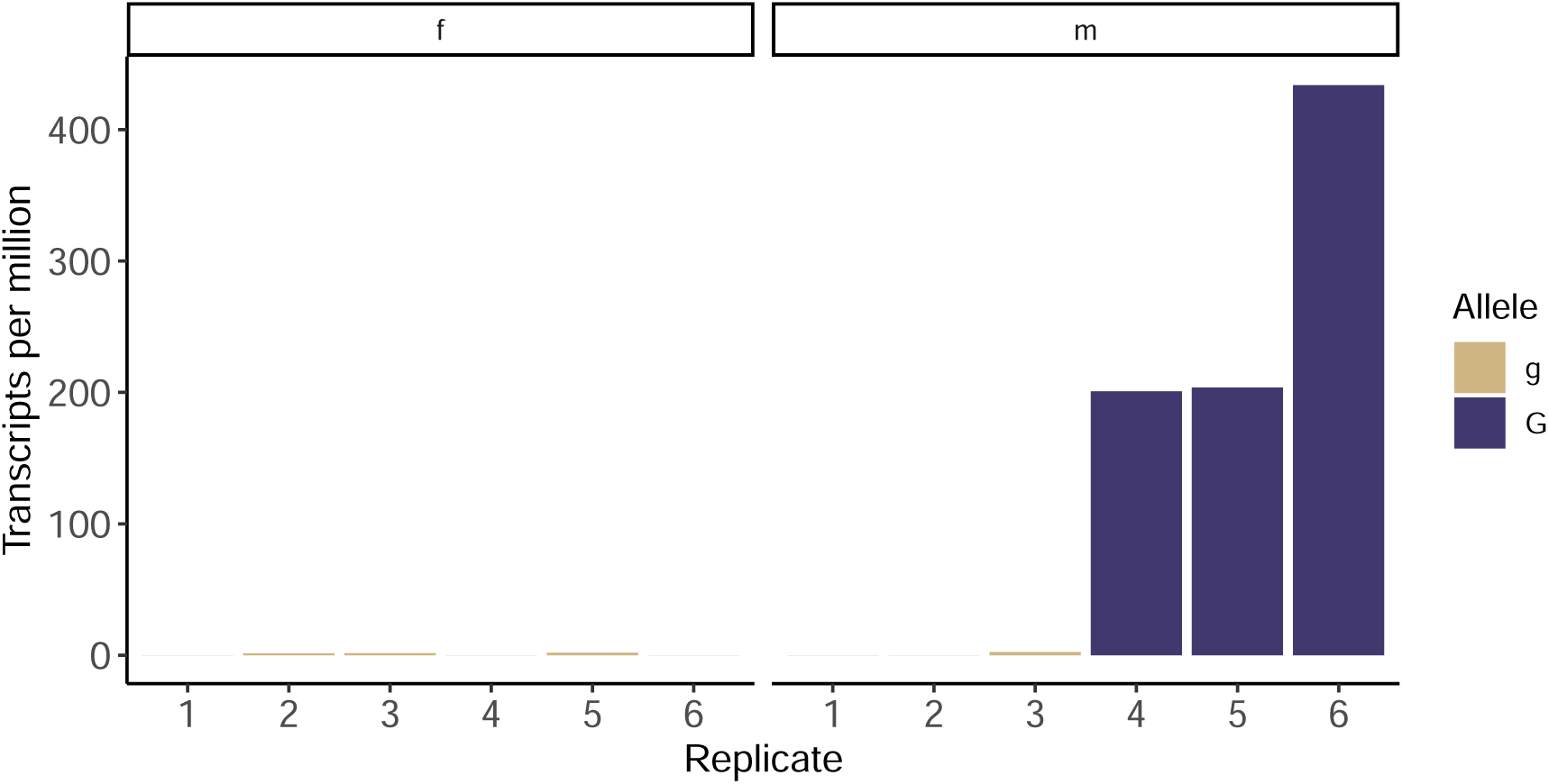
Reanalysis of RNA-seq experiment from Qin *et al*. (2021). The authors generated RNA-seq from male (m) and female (f) flower buds in the season prior to flowering. We measured relative read depth in each sample at SNPs within *TPPD-1* that are fixed between *J. mandshurica* G-locus haplotypes, ascertained from whole-genome resequencing data 60 individuals of this species (36 *G*, 84 *g* haplotypes) (Xu *et al*. 2021). Similar to the data from (Li *et al*. 2022b), these data indicate that *TPPD-1* is relatively highly expressed in male catkin buds, and that this is driven by allele-specific expression of the *G* haplotype copy of *TPPD-1*. We verified that this is not a mapping artefact caused by reference bias by mapping the reads to assemblies of both haplotypes. We note the authors described sampling both “male-precursor” and “female-precursor” types, which we take to mean protandrous and protogynous. Two of the male bud samples with high expression are labelled as “female-precursor,” while one is labelled “male-precursor.” These three samples contain multiple SNP alleles that are fixed in a large sample *J. mandshurica G* haplotypes, so we infer they are all from protogynous trees. Notably, RNA-seq reads from these samples contain only *G* variants and no *g* variants. While one explanation is that these individuals are homozygotes for the *G* allele, we saw zero *GG* heterozygotes in whole-genome resequencing of 60 individuals of this species, and relatedness estimates are consistent with these samples representing 3 unique genotypes, so this is extremely unlikely. Moreover, expression of the *g*copy is low in all samples and completely missing in several in both this dataset and in Fig. S10. Thus, we conclude this pattern reflects a strong bias in allele-specific expression toward the *G*haplotype copy of *TPPD-1* in male catkin buds of protogynous *J. mandshurica* individuals.

**Figure S12:**
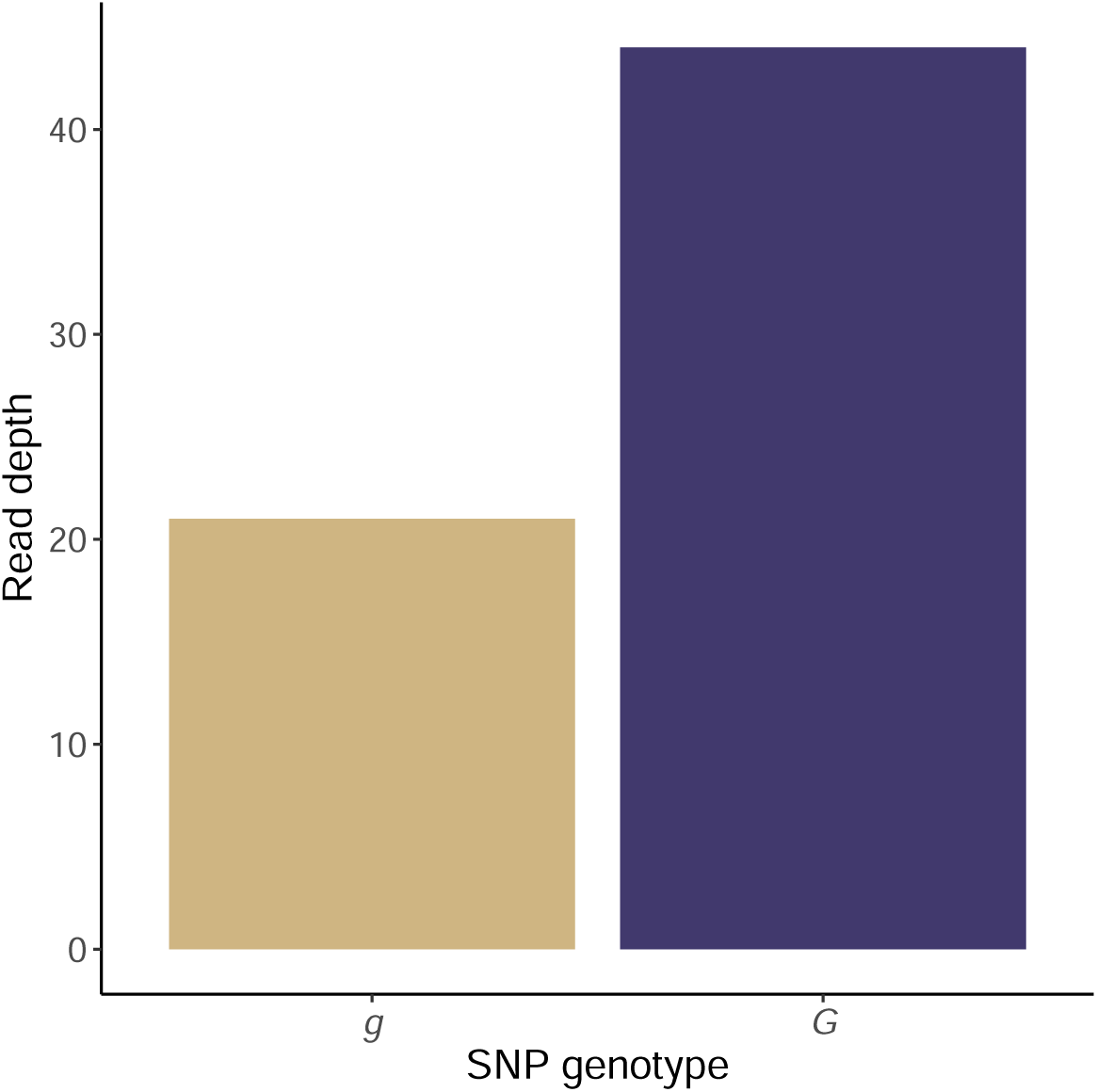
Reanalysis of RNA-seq data generated by Dang *et al*. (2016). The authors sequenced an RNA library pooled from four different tissues (leaf, bud, male flower, and female flower) from a single individual of *J. regia*. We identified this individual as a heterozygote by the presence of numerous SNPs in the *TPPD-1* coding region that we found to be fixed between G-locus haplotypes in a large sample (113 individuals) of this species. We measured allelic depth at 13 of these SNPs in coding sequence to quantify relative transcript abundances. The difference in read depth is statistically significant (P *<* 0.01). Shown for reads mapped to an assembly of the *g* haplotype. 11 out of 13 SNPs show the same direction of effect and with similar magnitude. Two SNPs show the opposite trend, both of these are in the last exon where divergence is higher, suggesting this could be an artefact of reference bias.

**Figure S13:**
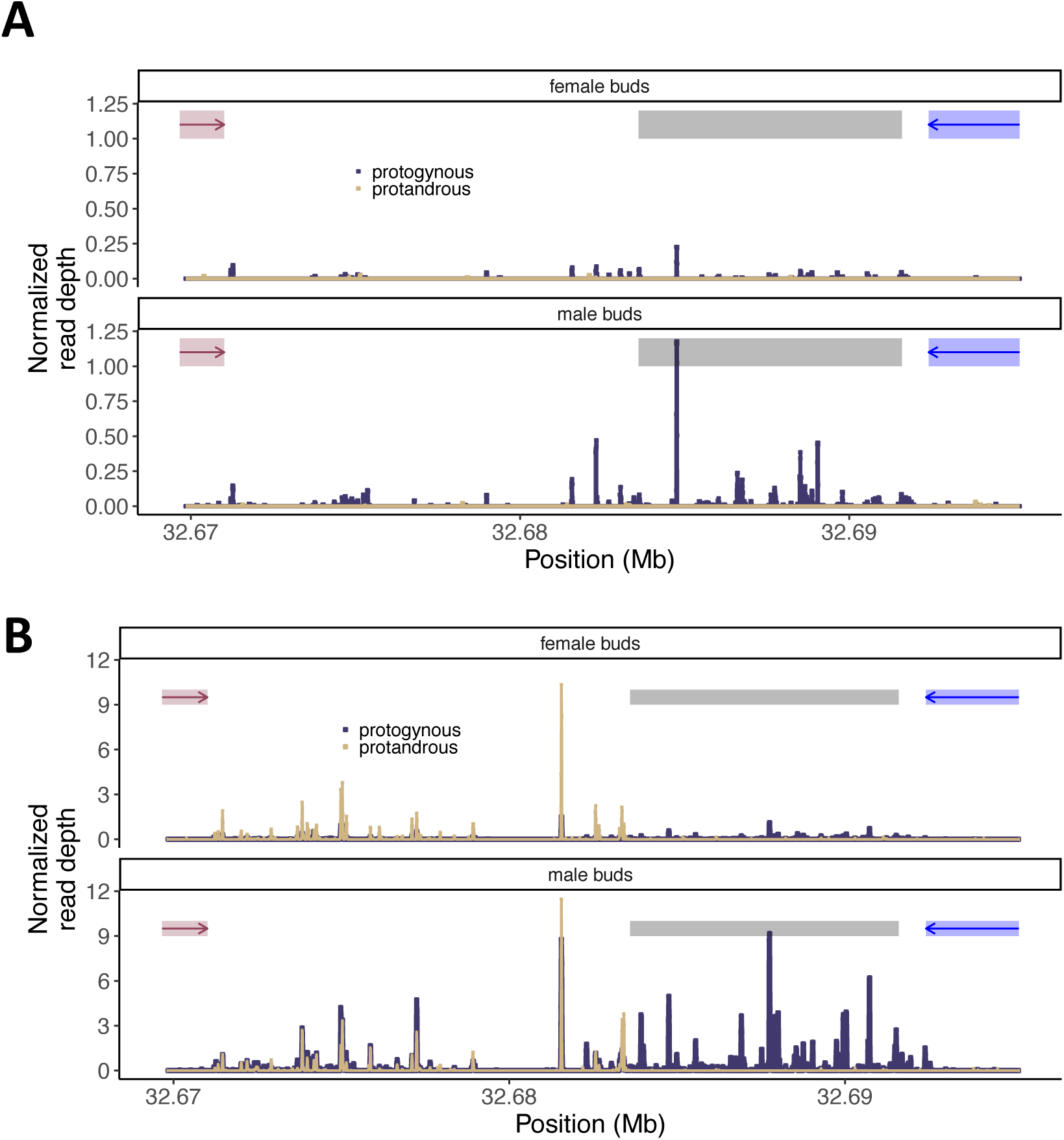
Reanalysis of a small RNA sequencing experiment from Li *et al*. (2023), where the authors sequenced small RNAs from male and female flower buds of *J. mandshurica*. **(A)** Small RNAs that map uniquely within the *G* haplotype of *J. mandshurica* with 1 bp mismatch tolerance. **(B)** Small RNAs that map to at most 10 locations in the genome with 1 bp mismatch tolerance. Colored bars indicate positions of bordering genes (*TPPD-1* shown in blue, *NDR1/HIN1* –like gene in red). Gray bar indicates the location of the *GJ1* indel.

**Figure S14:**
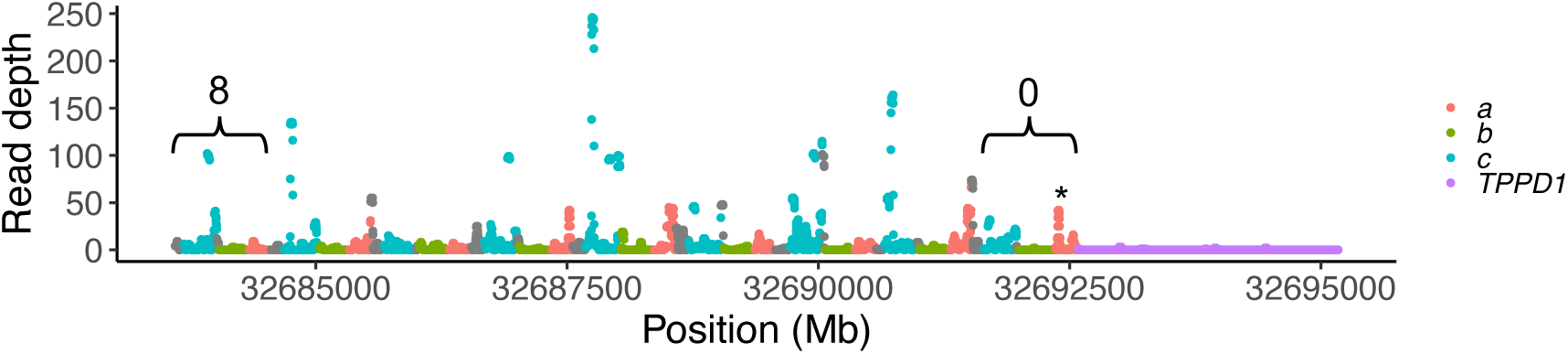
Sequencing depth of small RNAs from male bud tissue of a protogynous *J. mandshurica* individual, colored according to subunit of the *GJ1* indel. Repeat zero corresponds to the sequence found in both G-locus haplotypes just downstream of *TPPD-1* coding sequence. Subunit *a* of repeat zero (**\***) is the 3’ UTR of *TPPD1*. Total coverage shown across three technical replicates for reads that map to at most 10 locations in the genome, allowing for a single base pair mismatch. Data from (Li *et al*. 2023).

**Figure S15:**
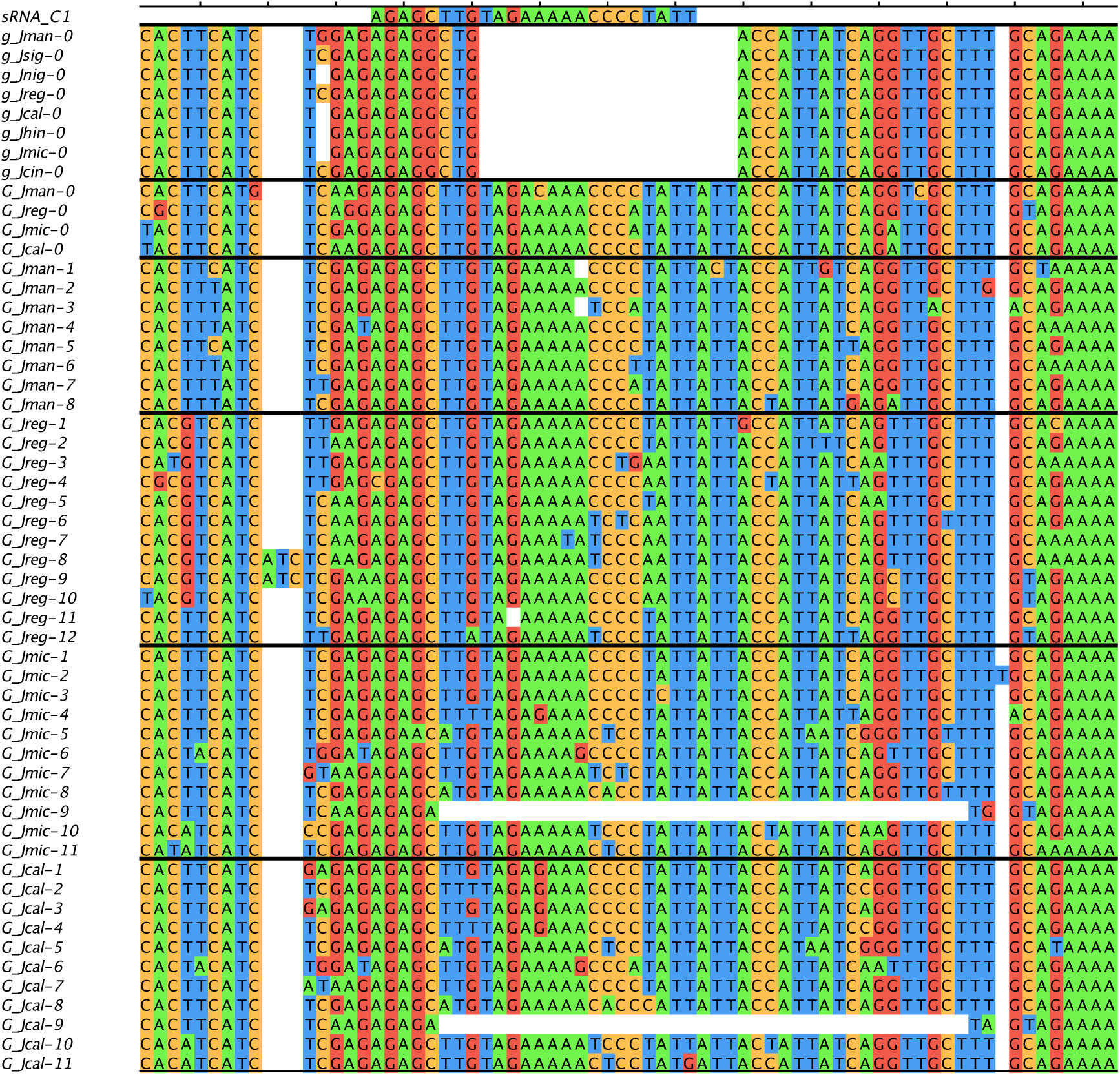
An abundant small RNA in male catkin buds of protogynous genotypes of *J. mandshurica*(sRNA C1, 24bp, top) maps to subunit *GJ1c* of the *GJ1* indel (see peak values in Fig. S13). We see conservation of this sequence within *GJ1c*repeats across species, and the sequence matches a site 800 bp downstream of *TPPD-1* 3’ UTR that is missing in *g* haplotypes. 3’ UTR sequences of *g* and *G* haplotypes are shown at top, and *GJ1* repeat sequences from *G* haplotypes of four *Juglans* species below.

**Figure S16:**
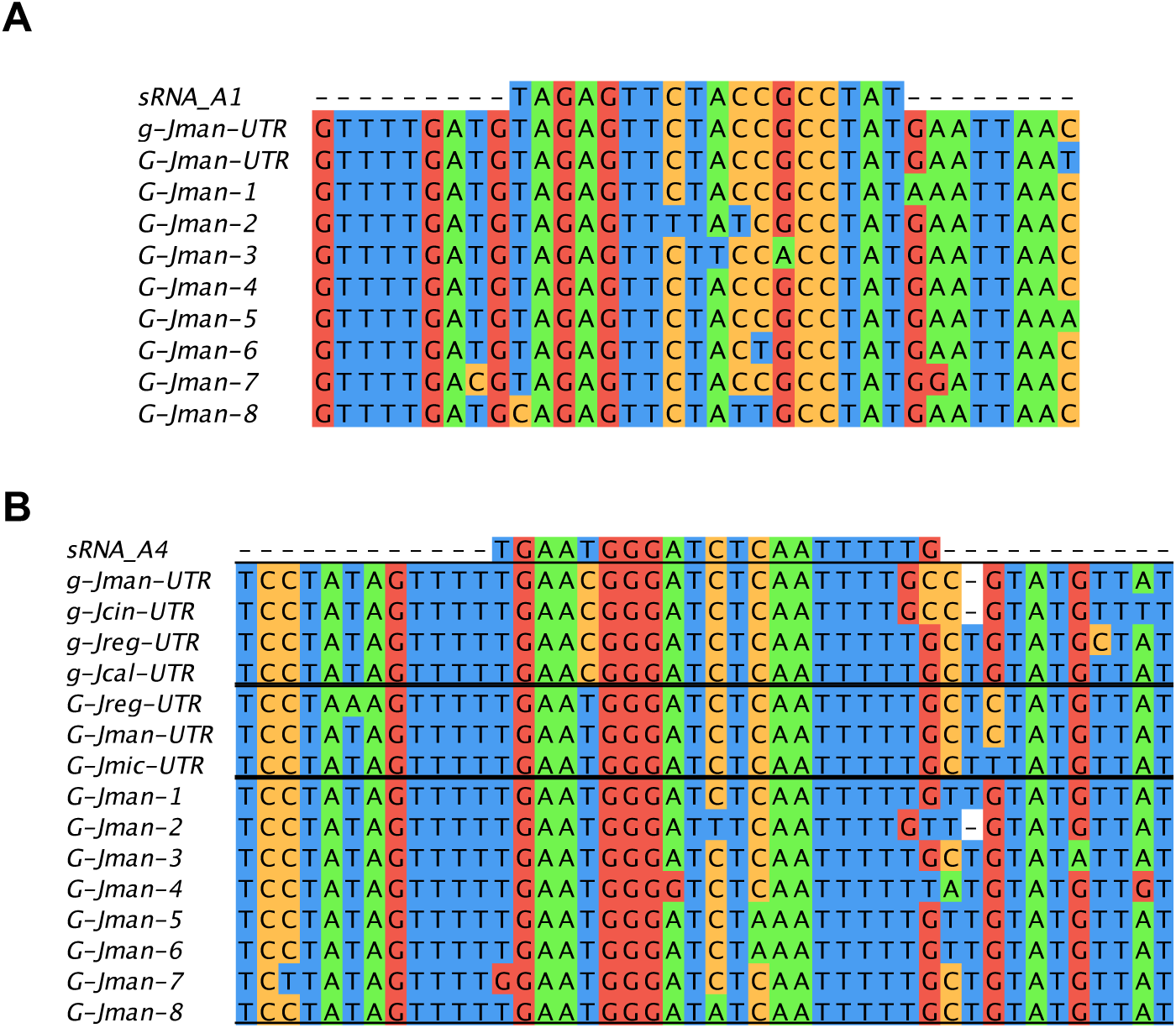
Several small RNAs found in catkin buds of protogynous genotypes of *J. mandshurica*match sequence in the 3’ UTR of *TPPD-1*. **A)** A small RNAs that matches both *G* and *g* 3’ UTR sequences. **B)** A small RNA that matches only *G* 3’ UTR sequence and overlaps a trans-species SNP polymorphism within the 3’ UTR.

**Figure S17:**
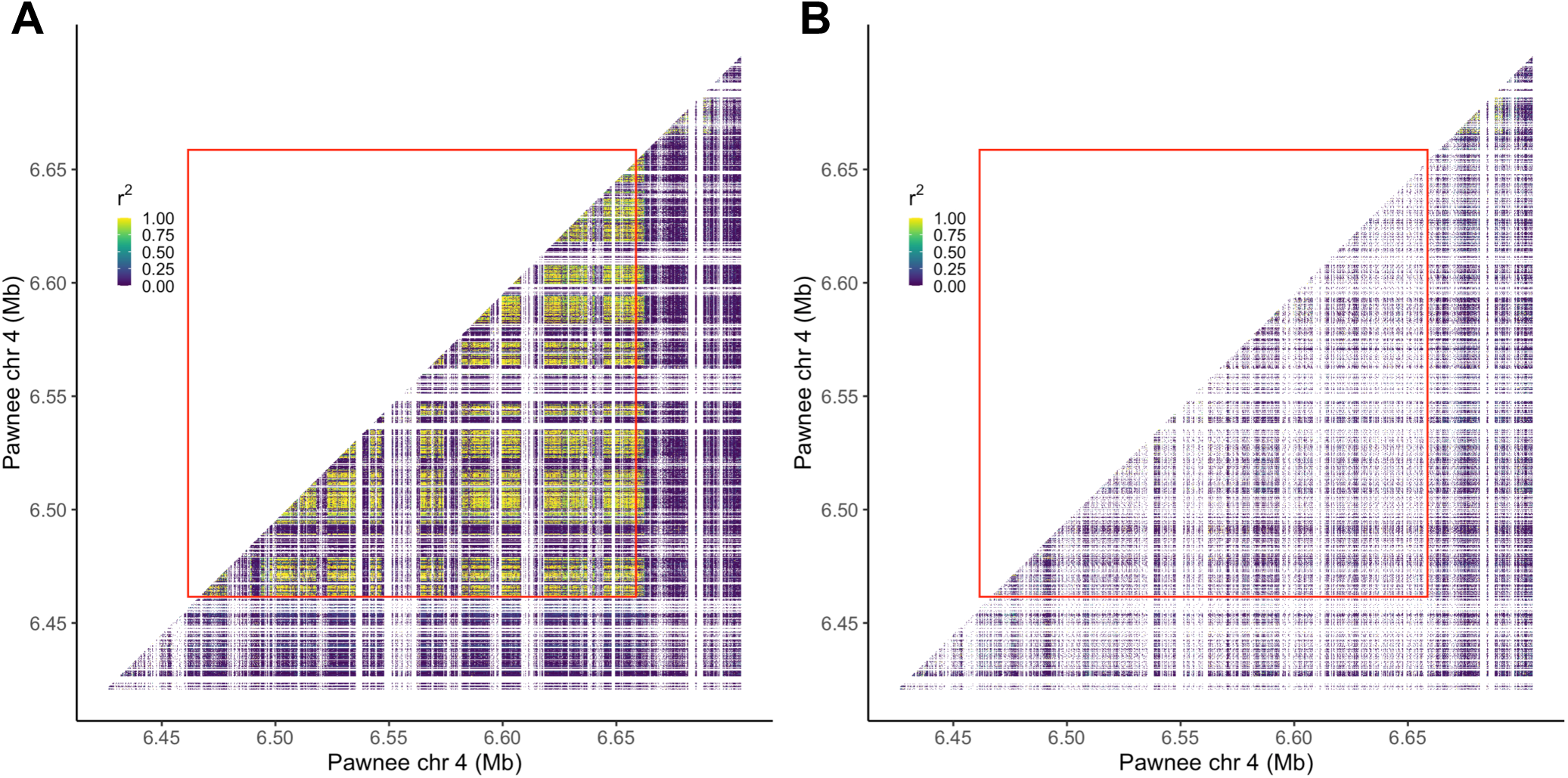
(**A**) Linkage disequilibrium is exceptionally high across the Carya G-locus, consistent with a lack of recombination between *G* and *g* haplotypes. Color indicates the strength of Pearson correlation between diploid genotypes at two loci. Red lines indicate the endpoints of genes that fall within the GWAS peak for dichogamy type in pecan. Shown for variant calls against the ‘Pawnee’ reference. **(B)** Using only genotypes of individuals homozygous for the recessive haplotype (*gg*), there is no signal of elevated LD across the same region, indicating recombination reduced recombination between *G* and *g* haplotypes but not between *g* haplotypes.

**Figure S18:**
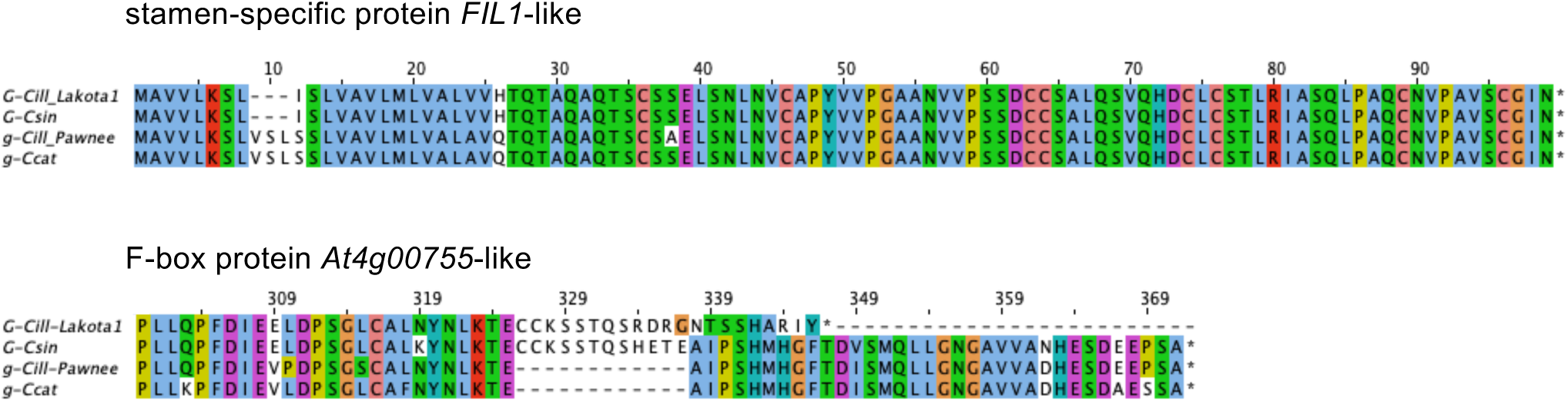
Conserved indels within coding sequence at the *Carya* G-locus. Top – candidate gene *FIL1* shows a three amino acid indel that is a shared difference between G-locus haplotypes across North American and East Asian clades of *Carya*. Bottom – F-box protein *At4g00755* –like shows evidence of a conserved coding indel near the terminus of the polypeptide.

**Figure S19:**
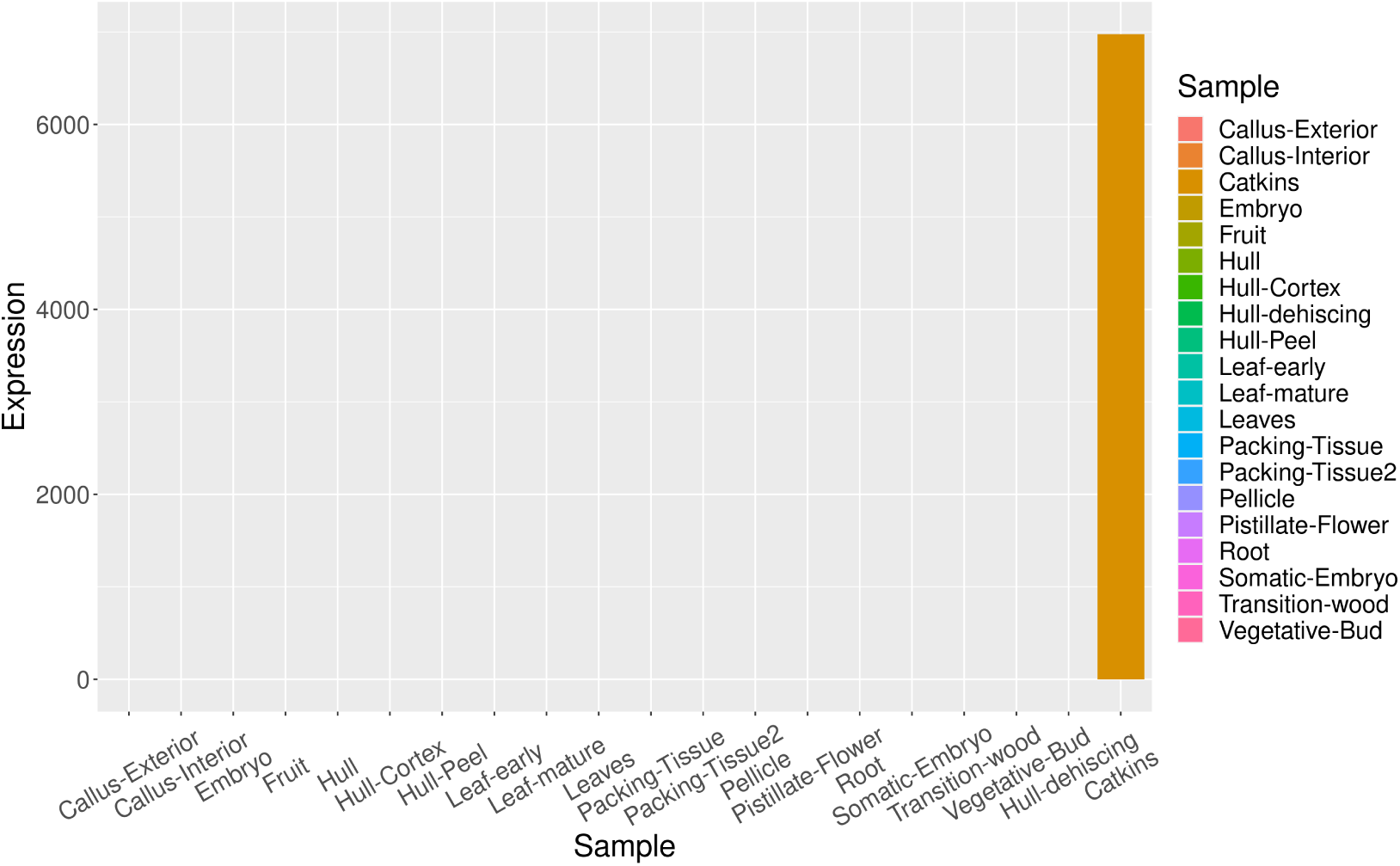
Gene expression levels of the *J. regia* ortholog of *FIL1* in 20 transcriptomic libraries from different tissues. Results from the literature indicate that *FIL1* –homologs are expressed uniquely or predominantly in stamens Nacken *et al*. (1991). Consistent with this, we detected abundant expression of *FIL1* in catkins of *J. regia*, and no trace of expression in other tissues. Y-axis measures Fragments Per Kilobase of transcript per Million mapped reads.

**Figure S20:**
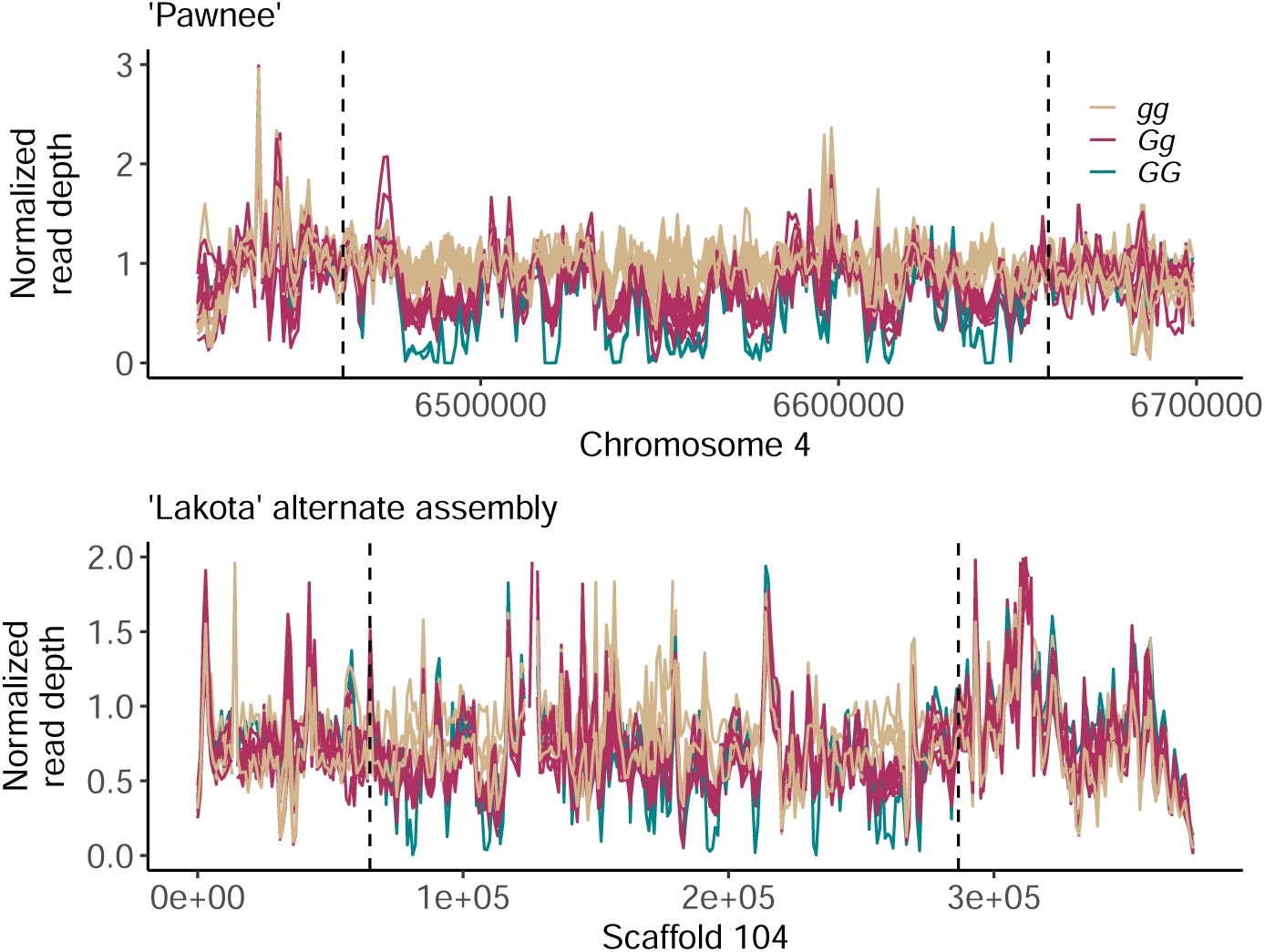
Normalized average read depth of pecan varieties in 1kb windows across the pecan G-locus for two assemblies of the *g* haplotype. **Top)** ‘Pawnee’ is protandrous (*gg*) so the assembly is known as the *g* haplotype a priori. **Bottom)** The alternate assembly of ‘Lakota’ (*Gg*) shows the same coverage pattern as for ‘Pawnee’, which is reversed compared to the ‘Lakota’ primary assembly (see Fig 3C in main text).

**Figure S21:**
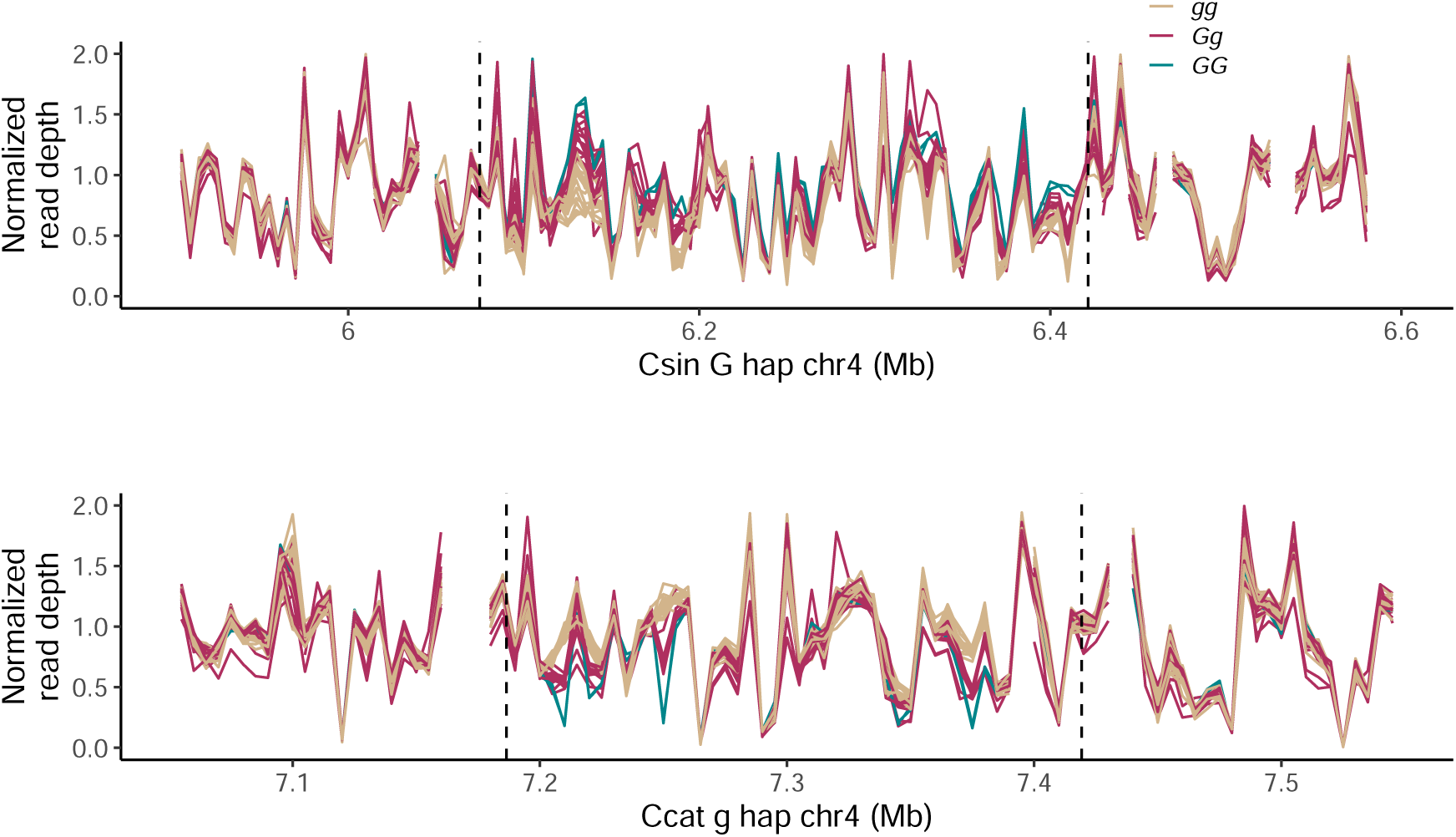
Read depth of pecan varieties (*C. illinoinensis*) of against two East Asian *Carya* genome assemblies. Top: Patterns of differential read depth by G-locus genotype indicate that the assembly of *C. sinensis* is of the *G* haplotype. The G-locus region in this assembly is roughly 350kb in length, closer in length to the pecan *G*haplotype than to the pecan *g* haplotype. Vertical indicate the outermost G-locus genes as defined in pecan, and these correspond roughly to the span of differential coverage patterns. Bottom: Differential read depth by G-locus genotype against the *C. cathayensis* assembly indicate it is an assembly of the *g* haplotype. The G-locus region in this assembly is roughly 230kb, closer in length to the *g* haplotype of pecan.

**Figure S22:**
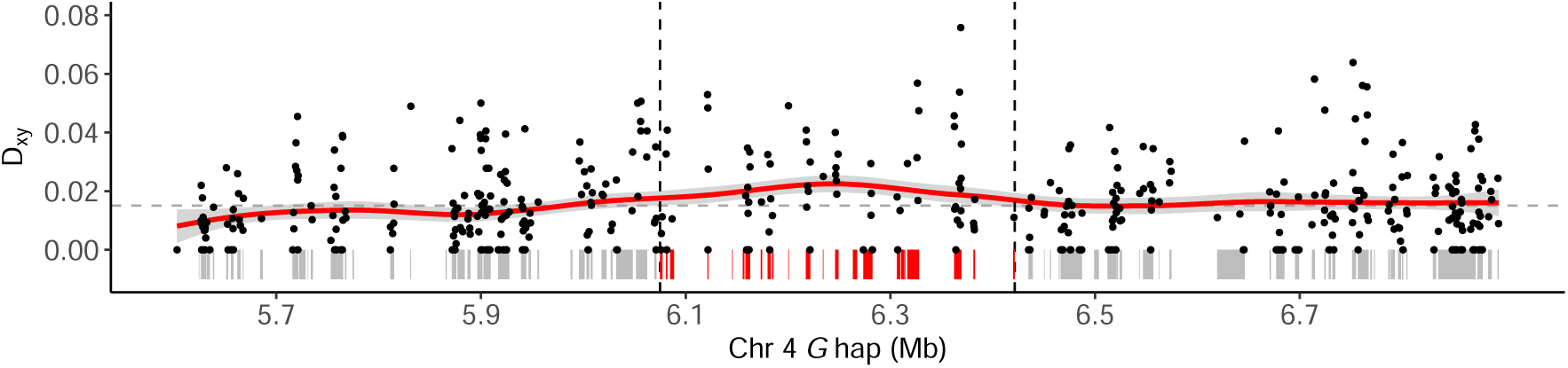
Nucleotide divergence in coding sequence between a genome assembly of *Carya sinensis* and one of *Carya cathayensis*. The elevated nucleotide divergence in the region syntenic with the pecan G-locus is further consistent with these two assemblies representing alternate haplotypes at the G-locus. Dotted horizontal line corresponds to the genome-wide weighted average for coding sequence. Dotted vertical lines indicate the positions of genes orthologous to those at the boundaries of the pecan G-locus. Below, shaded bars represent positions of genes. Red bars indicate orthologous genes contained within the region syntenic to the pecan G-locus.

**Figure S23:**
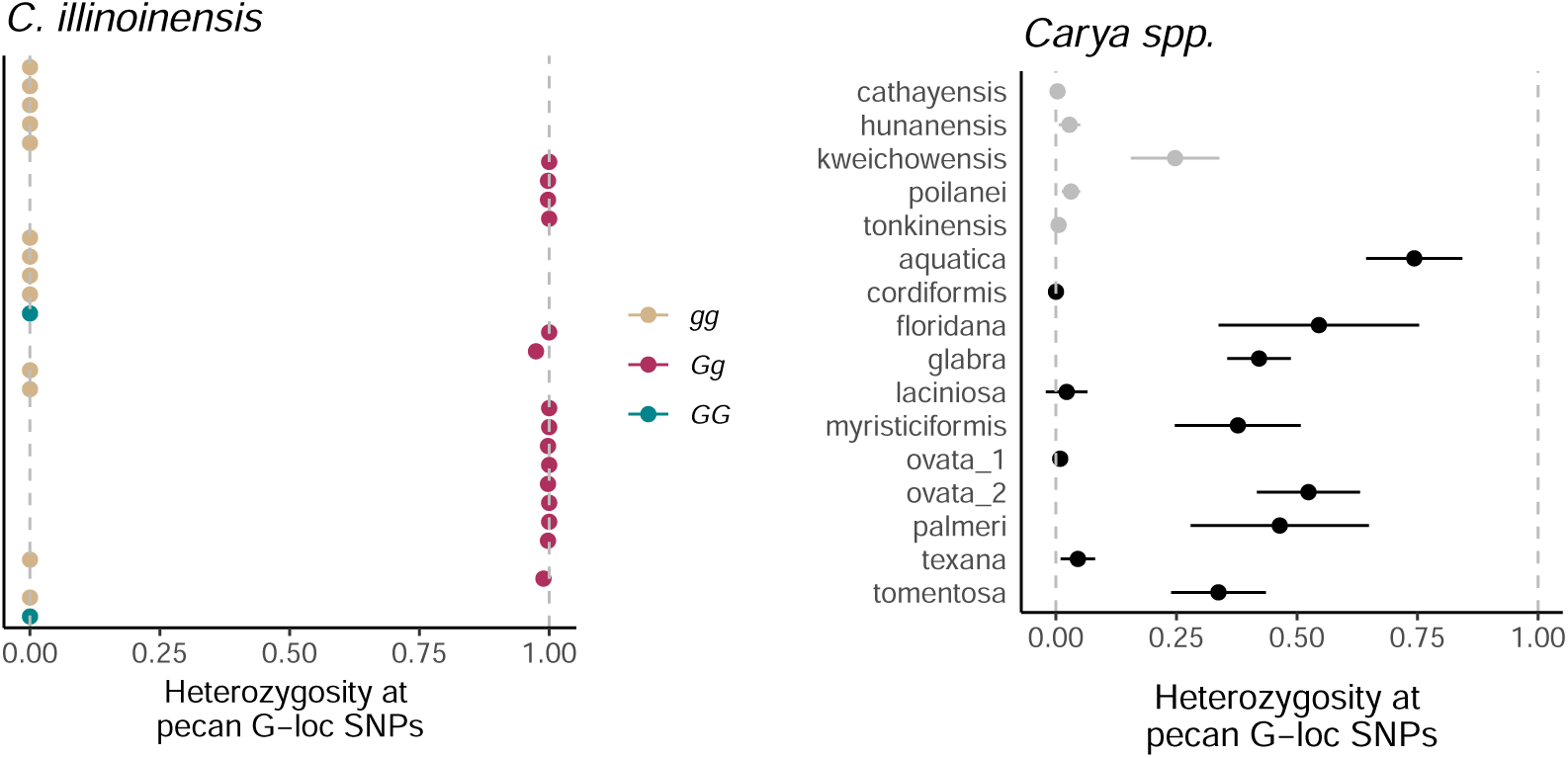
Individual-level heterozygosity at a set of 454 SNPs in G-locus coding sequence that were ascertained as fixed differences between *GG* and *gg* genotypes in pecan. **Left)**Heterozygosity at these SNPs is very close to 100% for heterozygotes for the G-locus haplotypes, confirming that these vast majority of SNPs are indeed fixed differences. **Right)** Heterozygosity at the same set of SNPs for 16 individuals representing 15 species of *Carya* spanning the divergence between North American (black) and East Asian (gray) clades. Error bars show 95% confidence intervals for the proportion of heterozygous SNPs. Note two individuals of *C. ovata*, one of which appears to be a heterozygote for the G-locus haplotypes and the other a homozygote. The putative heterozygote in the East Asian clade, *C. kweichowensis*, is heterozygous at a lower proportion of SNPs, which would be explained by many of these SNPs having arisen after the divergence between East Asian and North American clades. Consistent with this, we note that the individual with the highest proportion of heterozygous SNPs is from *C. aquatica*, which is the sister species to *C. illinoinensis*.

**Figure S24:**
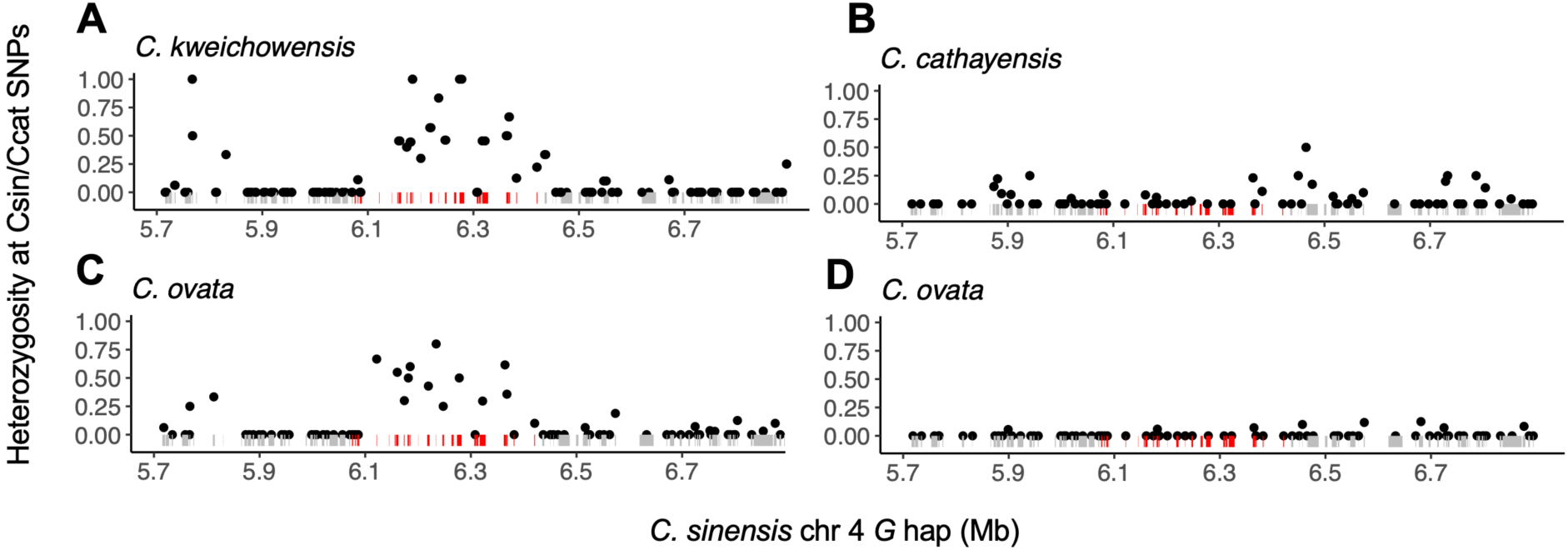
Individual-level heterozygosity at SNPs that distinguish the *Carya sinensis* (*G*) and *Carya cathayensis* (*g*) assemblies. Four individuals from Fig. S23 are shown to illustrate trans-species polymorphism in both North American and East Asian clades for G-locus SNPs ascertained from the East Asian *Carya* assemblies. Each point corresponds to a single gene. Below, bars indicate positions of predicted genes. Red bars correspond to genes orthologous to those associated with dichogamy type in pecan (*C. illinoinensis*). **A,B)** Two individuals from the East Asian clade of *Carya*, where (A) of which is evidently a heterozygote for the G-locus haplotypes and (B) is a homozygote (see also Fig. S23). **C,D)** Two individuals of *C. ovata* from the North American clade, where (C) is a heterozygote for G-locus haplotypes and (D) is a homozygote.

**Figure S25:**
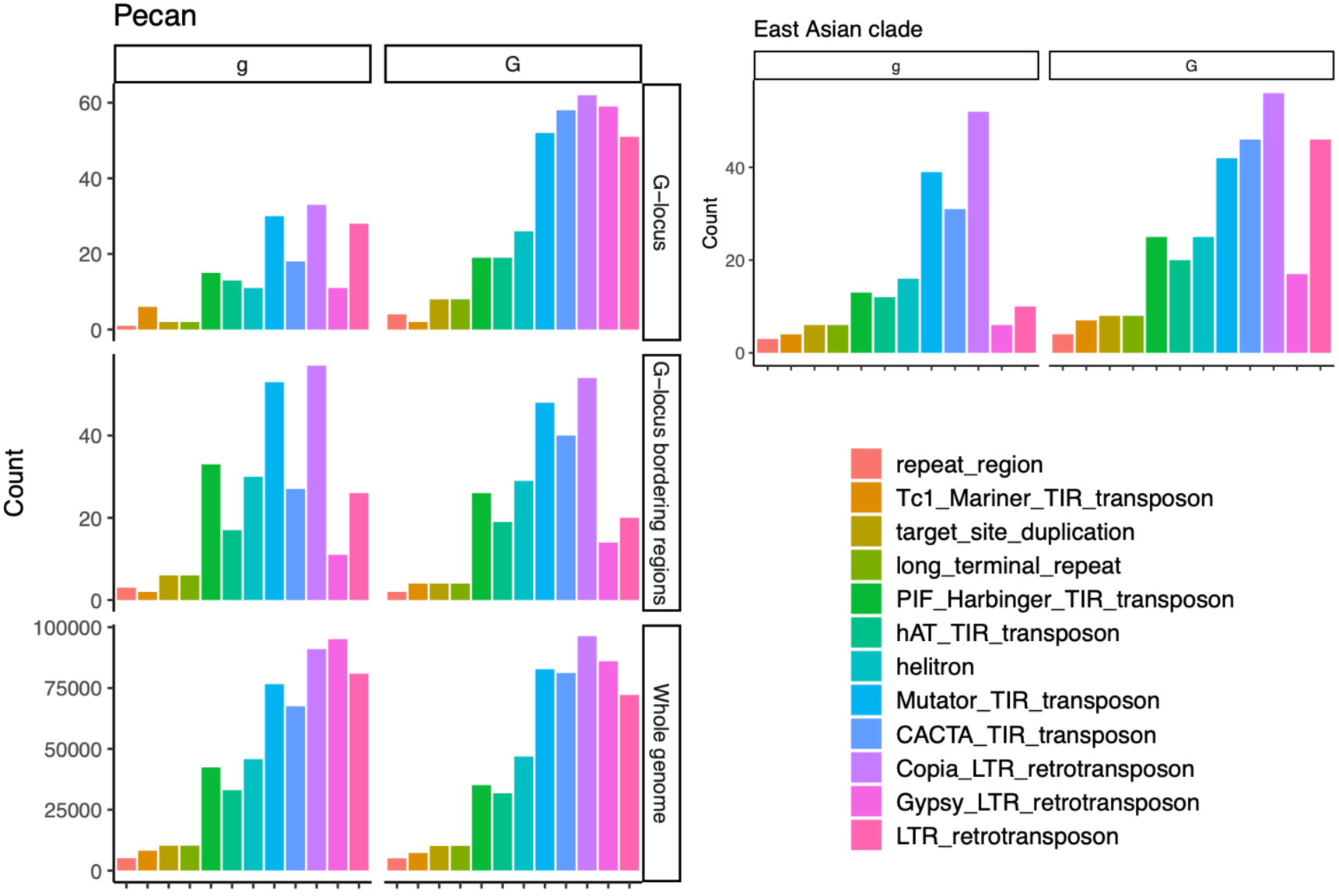
Transposable element (TE) content within *Carya*G-locus haplotypes. **(Left)** Number of TEs by category within the G-locus, in 150kb of flanking sequence on either end of the H-locus, and across the entire genome for two pecan (*C. illinoinensis*) assemblies containing alternate G-locus haplotypes. **(Right)** TEs within the G-locus for assemblies of haplotype within the East Asian clade. In this case, the assemblies are from different species, so for simplicity we omit showing background TE content outside of the G-locus, but note the relative increase in this region for *C. sinensis* compared to *C. cathayensis* diverges from the genome-wide average pattern.

**Figure S26:**
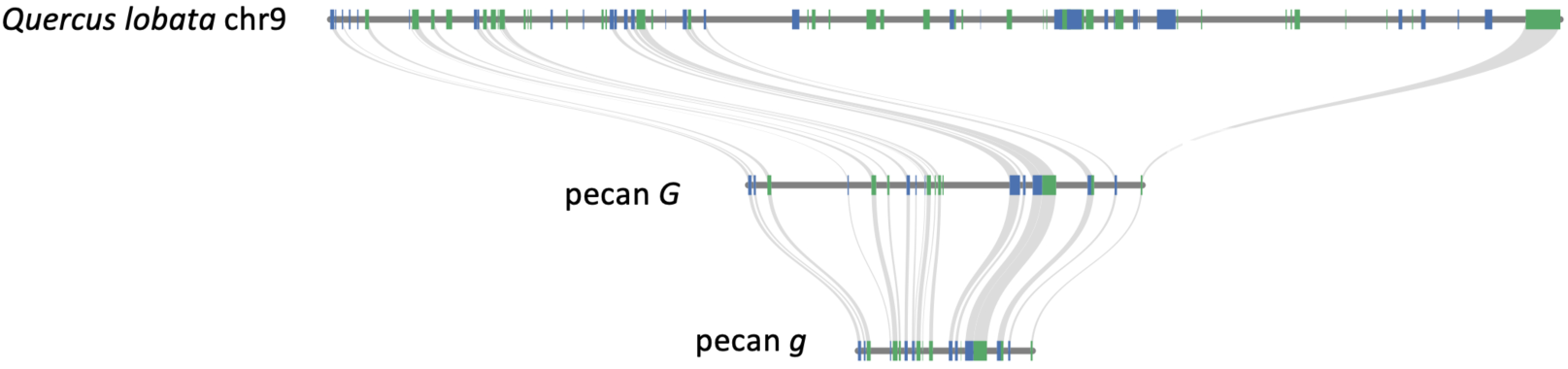
Gene-level synteny for pecan G-locus haplotypes and *Quercus lobata*. Gray bands connect orthologous genes. Genes are colored by strand (blue – plus, green – minus). Shown for the 20 predicted genes that fall within the pecan GWAS peak for dichogamy type and are present in the annotation of both pecan assemblies.

**Figure S27:**
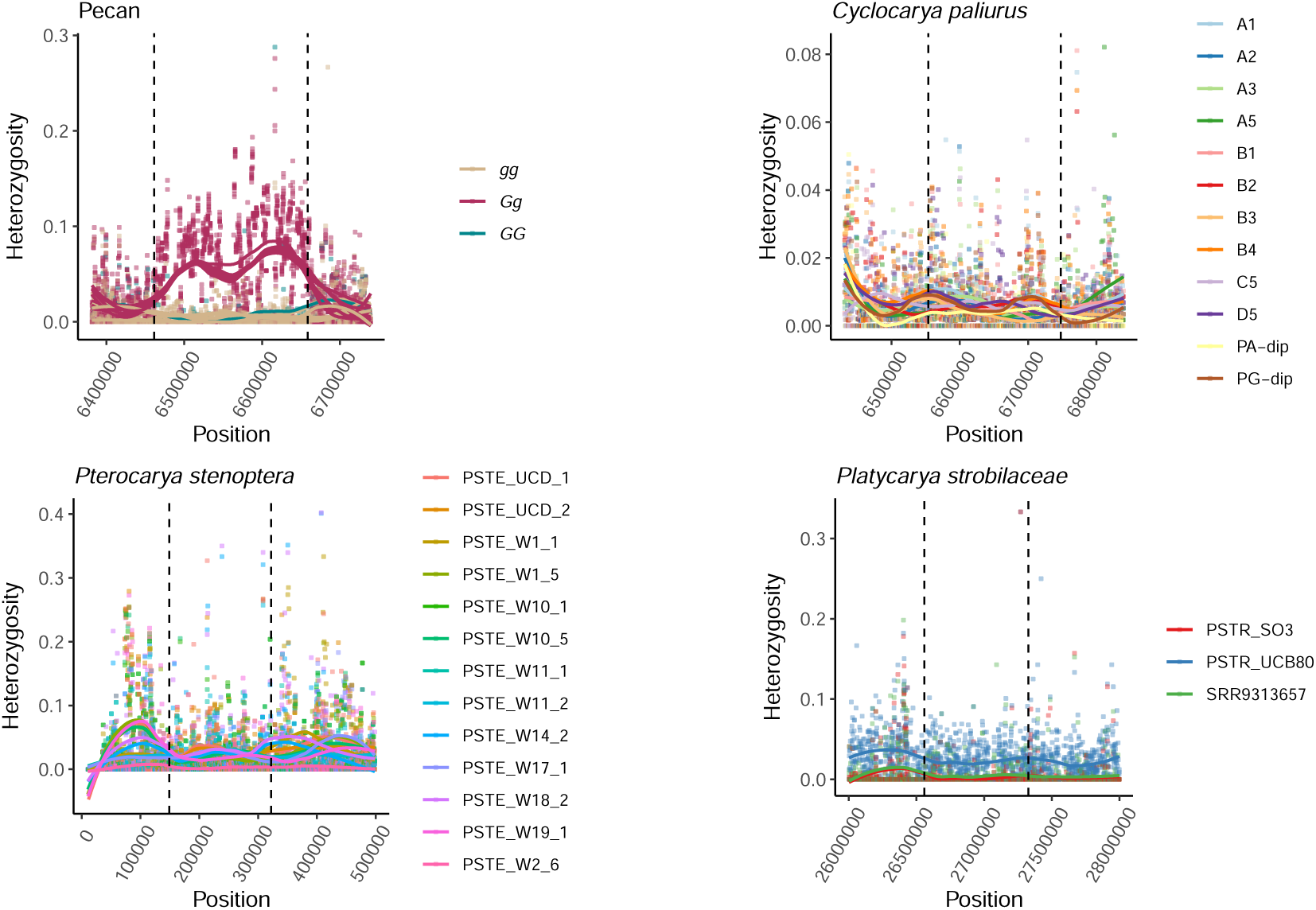
**Topeft**) Heterozygosity in 1kb windows for pecan varieties of known genotype at the *Carya* G-locus. **Topright)** Heterozygosity in 1kb windows for 12 diploid *Cyclocarya paliurus* individuals across the region syntenic with the *Carya* G-locus. Two of these individuals are of known dichogamy type (Qu *et al*. 2023). **Bottomleft)** Heterozygosity in 1kb windows for 13 diploid *Pterocarya stenoptera* individuals across the region syntenic with the *Carya* G-locus. **Bottomright)** Heterozygosity in 1kb windows for 3 diploid *Platycarya strobilaceae* individuals across the region syntenic with the *Carya* G-locus. Lines are LOESS smoothing curves fit for each individual. Dotted lines show the endpoints of the outermost G-locus orthologous genes in each assembly.

**Figure S28:**
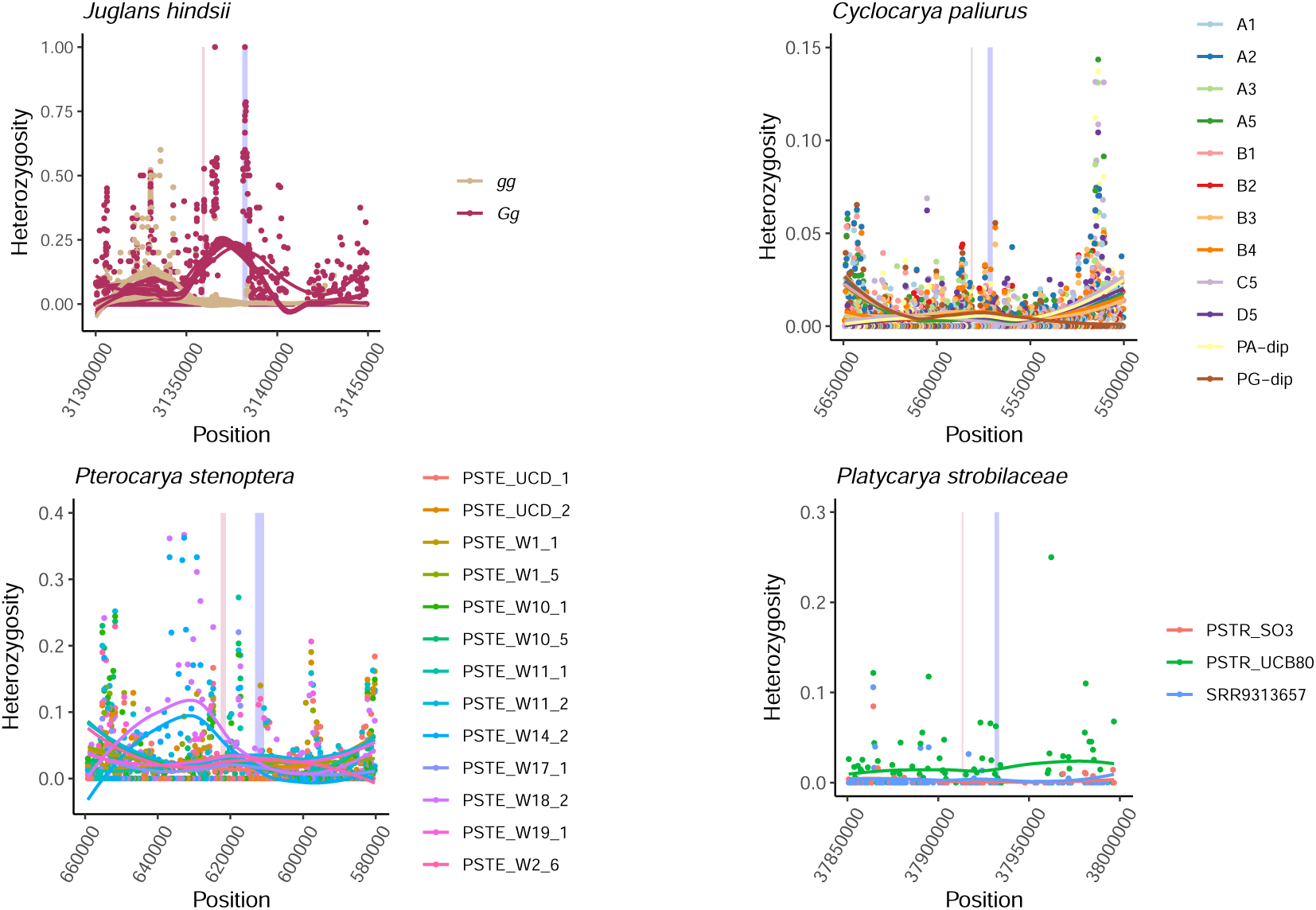
**Topleft**) Heterozygosity in 500bp windows for protandrous and protogynous individuals of Northern California black walnut (*J. hindsii* at the *Juglans* G-locus. Note the clear separation of heterozygotes from homozygotes. Lines are LOESS smoothing curves fit for each individual. Blue rectangle indicates the location of *TPPD-1*, red rectangle indicates the location of a *NDR1/HIN1* –like gene. **Topright)** Heterozygosity in 500bp windows for 12 diploid *Cyclocarya paliurus* individuals across the region syntenic with the *Juglans* G-locus. Two of these individuals represent each dichogamy type (Qu *et al*. 2023). **Bottomleft)** Heterozygosity in 500bp windows for 13 *Pterocarya stenoptera* individuals across the region syntenic with the *Juglans* G-locus. Here, we note two individuals appear highly heterozygous to the left hand side of the *NDR1/HIN1* –like gene, but that individuals also show high heterozygosity in the corresponding region in *J. hindsii* that is not associated with dichogamy type. We also note that we did not detect G-locus structural variants in *P. stenoptera* and *C. paliurus* (Fig. **??**), and our phylogeny inference (Fig. 2C) suggests we should not expect to find the *Juglans* G-locus structural variant segregating in *Pterocarya*. **Bottomright)** Heterozygosity in 500bp windows for 3 individuals of *Platycarya strobilaceae*, another heterodichogamous species in Juglandaceae, in the region syntenic with the *Juglans* G-locus.

## Notes

### Competing Interest Statement

The authors have declared no competing interest.

### Summary of Updates

Updated results to include analyses of gene expression of TPPD-1 and small RNA; minor updates to various analyses and figures.

